# Autophagy as a regulatory node integrating the carbon/nitrogen ratio with the auxin-driven chloronema-to-caulonema transition in *Physcomitrium patens*

**DOI:** 10.64898/2025.12.28.696759

**Authors:** Georgina Pettinari, Franco Liberatore, Veronica Mary, Martin Theumer, German Robert, Magdalena Bezanilla, Ramiro Lascano, Laura Saavedra

## Abstract

Plants, as sessile living systems, have the capacity to respond to both internal and external signals by modulating their growth and developmental programs. Autophagy is a recycling process which is known to respond to these signals, yet its role as an integrative regulatory node coordinating environmental inputs and developmental outputs remains underexplored, particularly outside model angiosperms like *Arabidopsis thaliana*. Here, we investigate how the fluctuating supply of carbon and nitrogen modulates autophagy in relation to the auxin-driven chloronema-to-caulonema transition in the bryophyte *Physcomitrium patens*. We show that autophagy-deficient mutants display a loss of coordinated plasticity in response to the changing C/N supply ratio, exhibiting an initially enhanced yet ultimately unsustainable response to nitrogen deficiency alongside attenuated responses to external sucrose. These altered responses reflect a modified basal state in the mutants, evidenced by higher intrinsic caulonemal growth, sucrose accumulation, altered auxin homeostasis, and differential auxin-related gene expression. Consistent with this, exogenous auxin elicits a diminished developmental response but accelerates senescence in the mutants. Finally, we demonstrate how autophagic flux is dynamically modulated by the C/N supply ratio and show that it is activated when this ratio is most unbalanced. We propose that high autophagic activity is associated with chloroplast-rich chloronemal cells, whereas caulonemata-inducing conditions decrease autophagic flux. This implies a cell-type-specific modulation of autophagy, where it is negatively correlated with caulonemal development. Together, our findings identify autophagy as a key regulator within the system that modulates ordered and adaptive responses to both external nutrient availability and internal hormonal signaling.

**HIGHLIGHTS:** Autophagy coordinates carbon/nitrogen supply with developmental programs in *P. patens*.

Autophagy negatively correlates with auxin-driven caulonemal development.

Autophagy-deficient mutants lose coordinated plasticity.

*atg* mutants exhibit a modified basal state with altered sugar and auxin profiles.

## 1. INTRODUCTION

As sessile organisms, plants continuously sense their environment and adjust their metabolism, physiology, morphology, growth and development to cope with changing conditions. This phenotypic plasticity is a capacity of the living system which allows plants to survive and thrive throughout their life cycle. To achieve this, cells remove damaged proteins, organelles and other debris that could cause malfunction, while recycling and replenishing nutrients to sustain their growth. Therefore, quality control systems, such as the ubiquitin/proteasome pathway and the autophagic route, are crucial for maintaining homeostasis and survival of plant cells. Whereas the ubiquitin/proteasome pathway specializes in small, short-lived proteins, autophagy is the primary degradative pathway for larger cytoplasmic material, such as organelles or protein aggregates (Dikic, 2017; Gross et al., 2025; Marshall & Vierstra, 2018; Petersen et al., 2024).

Autophagic induction in response to different stress conditions has been widely reported in plants (Avin-Wittenberg, 2019; Bassham et al., 2006; Filomeni et al., 2015; Jung et al., 2017; Kroemer et al., 2010; Lin L.-Y. et al., 2023; Liu Y. et al., 2009; Rose et al., 2006; Signorelli et al., 2019; Slavikova et al., 2008; Thirumalaikumar et al., 2021). Autophagy is also activated during cellular reprogramming triggered by different stimuli, like wounding or phytohormone application, removing cellular components that are no longer needed to ensure the correct execution of the new cell program (Kanne et al., 2022; Rodriguez et al., 2020). Beyond stress-induced responses, autophagy also plays essential roles in normal plant growth and development, including maintaining energy availability during the night and reallocating nutrients during senescence (Avila-Ospina et al., 2014; Chen et al., 2019; Guiboileau et al., 2012; Izumi et al., 2013; Li et al., 2015; Wang Y. et al., 2013). In addition, autophagy is involved in programmed cell death in plants, contributing to developmental processes such as xylem maturation, as well as modulating the hypersensitive response to incompatible pathogens (Courtois-Moreau et al., 2009; Hofius et al., 2009; Liu Y. et al., 2005; Üstün et al., 2017).

Most of our knowledge of autophagy comes from studies on a very limited number of angiosperms; however, its integration with hormone signaling and nutritional stress remains poorly understood in other plant lineages. In this context, bryophytes are particularly valuable to study cellular processes involved in stress responses and homeostasis maintenance since they were the first group to successfully colonize the xeric, nutrient-poor terrestrial environment. Land plants have developed the complex stress responses needed to survive in this fluctuating environment, and most of these are present in bryophytes (Dong et al., 2025; Kürschner, 2004; McDaniel, 2021; Wang Q.-H. et al., 2022; Sakil, 2025). Studying this acquired plasticity in bryophytes allows us to understand how these response systems, such as autophagy, coordinate stress signaling with growth and developmental programs. Their relatively simple morphology and life cycle make them ideal to study such processes. Among bryophytes, the moss *Physcomitrium patens* has been widely established as a model organism for studying evolutionary plant cell biology and developmental processes (de Keijzer et al., 2021; Naramoto et al., 2022; Prigge & Bezanilla, 2010; Rensing et al., 2020).

The life cycle of *P. patens* is predominantly haploid and begins with the germination of a spore to form a chloronemal initial cell. This cell divides to produce chloronemal filaments (chloronemata), which can differentiate into caulonemal cells. These cells can further divide and form caulonemal filaments (caulonemata), from which secondary chloronemal or caulonemal branches can be produced. Collectively, chloronemata and caulonemata are known as protonemata. Three-dimensional buds later emerge from caulonemal filaments and give rise to leafy gametophores (Jaeger & Moody, 2021; Rensing et al., 2020). Chloronemal and caulonemal cells represent two different phases of *P. patens* filamentous growth. Both are tip-growing cells, yet they are morphologically and functionally distinct (Menand et al., 2007b; Vidali & Bezanilla, 2012). Chloronemal cells can be considered source tissues, analogous to the leaves of vascular plants, since they are rich in chloroplasts and actively perform photosynthesis. In contrast, caulonemal cells are longer, faster-growing cells with fewer and less developed chloroplasts, and colonize the substrate. Thus, similarly to the root hairs of vascular plants, caulonemata are responsible for substrate exploration and nutrient acquisition.

The transition from chloronemata to caulonemata is tightly regulated by auxins and depends not only on indole-3-acetic acid (IAA) accumulation but also on proper auxin sensing and signaling (Bascom et al., 2025; Nemec-Venza et al., 2022; Thelander et al., 2018, 2019). In *P. patens*, auxin biosynthesis depends mainly on the IPyA (indole-3-pyruvic acid) pathway, where PpSHI/STY transcription factors and YUCCA flavine monooxygenases (PpYUCs) play a major role (Eklund et al., 2010a, 2010b; Landberg et al., 2021; Thelander et al., 2022). Polar auxin transport is regulated by PpPIN exporters, with PpPINA encoding the most abundant isoform, which transports auxin toward the tip of the protonemal filaments (Bennett et al., 2014; Fujita et al., 2008; Lüth et al., 2023; Viaene et al., 2014). Auxin-induced caulonemata development is regulated by the bHLH transcription factors PpRSL, which have a conserved function in regulating the development of tip-growing cells involved in nutrient acquisition and rooting functions in other plants (Jang et al., 2011; Jang & Dolan, 2011; Menand et al., 2007a; Pires et al., 2013). Caulonemal differentiation can also be induced by environmental factors such as light quality and nutrient availability. While nutrient-rich conditions and low light intensity favor chloronemata development, the transition to caulonemata is promoted when the plant faces nutritional deficiency (e.g., nitrogen or phosphorus starvation), or conditions of “high energy” availability (e.g., high light intensity, glucose/sucrose supplementation, or elevated CO_2_) (Mohanasundaram et al., 2024; Thelander et al., 2005). Interestingly, all of these conditions cause carbon/nitrogen (C/N) imbalances, which in turn modulate autophagic activity. This raises the so far unexplored hypothesis that autophagy may act as a regulatory node coordinating the fluctuating C/N supply with the auxin-driven chloronema-to-caulonema developmental transition.

Previous studies from our group demonstrated that the autophagic-deficient *atg5* and *atg7* mutants senesce prematurely, displaying hypersensitivity to nutritional starvation, as well as accumulating salicylic acid (SA) and IAA. They were shown to prioritize plant spread through filamentous growth at the expense of reduced three-dimensional (3D) growth, thereby delaying the transition to the adult phase of the moss life cycle (Pettinari et al., 2022). We also showed that autophagy is induced in protonemata in response to nitrogen deficiency as well as darkness. In the present work, we explore the relationship between nutrient status and autophagy, focusing on the regulation of a key step in the life cycle of *P. patens*, the chloronema-to-caulonema transition. We further characterize *atg* mutants, highlighting their reduced plasticity when faced with fluctuating carbon and nitrogen conditions, i.e., the C/N supply ratio. They display an altered basal state, evident in their higher intrinsic caulonemal growth and modified sugar and auxin homeostasis. We generated an additional *atg* mutant, *atg14*, which consistently shows an intermediate phenotype compared to *atg5*. Finally, we demonstrate how autophagic flux is modulated by the C/N supply ratio, and is activated when this ratio is most unbalanced. We propose that high autophagic activity is associated with chloronemal cells, whereas caulonemata-inducing conditions decrease autophagic flux, implying a negative correlation between autophagy and caulonemal development. Our findings suggest that autophagy acts as a key regulator, coordinating ordered and adaptive responses with external nutrient availability and internal hormonal signaling.

## 2. METHODS

### 2.1. Plant material and growth conditions

All experiments described in this study were performed with *Physcomitrium (Physcomitrella) patens ssp. patens* (Hedwig) ecotype ‘Gransden 2004’. Plant cultures were grown axenically at 24°C under a long-day photoperiod (16-h light/8-h dark) with a photon flux of 60-80 mmol.m^2^.s^-1^. *P. patens* protonemal tissue from WT (wild type), *atg5*, *atg7* and *atg14* lines was subcultured routinely at 7-day intervals on cellophane disks (AA packaging) overlaying Petri dishes (90 mm in diameter) that contained complete medium: BCDAT [0.92 g/L di-ammonium tartrate (C_4_H_12_N_2_O_6_), 0.25 g/L MgSO_4_.7H_2_O, 1.01 g/L KNO_3_, 0.0125 g/L FeSO_4_.7H_2_O, 0.25 g/L KH_2_PO_4_ (pH 6.5), 0.147 g/L CaCl_2_.2H_2_O, and 0.001% Trace Element Solution (0.055 g/L CuSO_4_.5H2O, 0.055 g/L ZnSO_4_.7H_2_O, 0.614 g/L H_3_BO_3_, 0.389 g/L MnCl_2_.4H2O, 0.055 g/L CoCl_2_.6H_2_O, 0.028 g/L KI, 0.025 g/L Na_2_MoO_4_.2H_2_O)] and 7 g/L agar.

Life cycle progression of the WT, *atg14* and *atg5* lines was observed using a microfluidic device with the help of Dr. Magdalena Bezanilla (Dartmouth College, Hanover, NH, USA). Homogenized protonemal tissue from WT, *Ppatg14-16*, and *Ppatg5* lines was loaded into the microfluidic chamber and incubated with Hoagland solution (Sigma-Aldrich, equivalent to liquid complete medium) under continuous light for 21 days. The medium was then replaced with Hoagland solution supplemented with 1 µM NAA, and the tissue was incubated for an additional 15 days under these conditions. Tissue was imaged after 1, 5, 10, and 15 days of treatment using a Nikon Ti microscope equipped with a 20× objective (NA 0.8) and a Nikon DS-Vi1 color camera. Images were acquired every 20 min over a 24-h period, with the tissue continuously illuminated with white light.

### 2.2 Plant growth assay

For external sugar response experiments, moss plants were grown from 1 mm^2^ spot cultures, obtained from 7-day protonema grown in complete medium, on cellophane overlaid ammonium-free medium (-NH_4_^+^, +NO_3_^-^, complete medium lacking di-ammonium tartrate), as well as ammonium-free media with added glucose, sucrose or mannitol, at 1%, 2,5% or 5% (w/v). For each genotype (WT, *atg5* and *atg7*) three inocula were placed per Petri dish, and three independent dishes were used per treatment. Plants were grown for 19 days and imaged at 7 time points (days 0, 5, 7, 10, 12, 14, and 19) using a GelDoc Go Gel Imaging System (BioRad). The images were processed using the scikit-image Python library and plant area was quantified. Representative day-19 plant images were acquired using an Olympus SZX16 stereomicroscope equipped with a DP71 color camera. Plant area over time was modelled using the following logistic growth function:

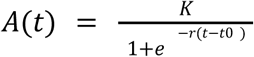

Where *A(t)* represents plant area at time *t*, *K* is the carrying capacity (maximum asymptotic area), *r* is the intrinsic growth rate, and *t*₀ is the inflection point of the curve. Models were fitted using nonlinear least squares regression. *K* and *r* were estimated independently for each genotype x medium combination, and *t*₀ was estimated separately for each environment. Nested models were compared using likelihood ratio tests and Akaike’s Information Criterion (AIC). Model assumptions were verified by visual inspection of residuals. Parameter contrasts between each treatment and its corresponding control (-NH_4_^+^) within each genotype were performed using Wald tests derived from the variance-covariance matrix of the fitted model. Confidence intervals were calculated as the estimate ± 1.96 × SE. All analyses were conducted in R (version 4.4.1) using the base stats and MASS packages.

### 2.3 Quantification of soluble sugars via HPLC

Protonemata of the WT, *atg5* and *atg7* genotypes were grown as previously described using ammonium-free medium medium for 7 days. Tissue was harvested at the end of the light period of day 7, and was ground in liquid nitrogen in microtubes. 80% cold ethanol was added in a w/v relation of 90 mg/750 μl, after which samples were incubated at 80°C for 30 min and then centrifuged at 14000 RPM for 10 min. The upper phase was collected, and one volume of pure acetonitrile was slowly added while on low shaking, after which the samples were left for 30 min at room temperature. Then samples were centrifuged at 14000 RPM for 15 min, the upper phase was collected and incubated at 90°C until evaporation. 500 μl of 75% acetonitrile was added, and samples were vortexed for 10 min. After that, samples were filtered using PDVF filters, and 10 μl were injected into an HPLC system. A Zorbax NH_2_ column was used (4,6 mm x 250 mm x 5 μm) with an optimized flux of 1,5 μl/min, each sample being run for 30 min using a mobile phase of acetonitrile/water 70/30%. Times of retention were: fructose 4,6 min; glucose 5 min; sucrose 6,5 min. Results are shown as the area under the curve (AUC) relative to fresh weight (FW).

### 2.4 Generation and identification of Ppatg14 knock-out mutants

The *atg14-ko* vector for generating mutants by homologous recombination was constructed by cloning approximately 1000 bp from the 5’ and 3’ ends of the *PpATG14* gene into a pUBW302 plasmid with G418 resistance. The resulting vector was linearized using the restriction enzymes XhoI and XbaI and introduced into WT by polyethylene glycol–mediated protoplast transformation according to Saavedra et al., 2015. Briefly, 6-day-old protonemal tissue from WT was incubated under gentle agitation (approximately 20 RPM) with 1% (w/v) Driselase (Sigma-Aldrich) dissolved in 8.5% (w/v) mannitol for 45 min at room temperature. The resulting protoplasts were filtered using a 100-µm mesh filter and washed three times with 8.5% (w/v) mannitol by centrifugation at 600 rpm. For each transformation, 0.3 ml of protoplasts at a density of 1.6 × 10⁶ protoplasts/ml, 10–15 µg of linear DNA from the plasmid of interest, and 0.3 ml of PEG solution (8.5% [w/v] mannitol, 1% [v/v] 1 M Ca(NO_3_)_2_, 0.1% [v/v] 1 M Tris-Cl pH 8.0, and 4 g PEG 4000) were used. Transformed protoplasts were then plated on Petri dishes containing PMRB medium (0.5 g/L di-ammonium tartrate (NH_4_)_2_C_4_H_4_O_6_, 0.25 g/L MgSO_4_.7H_2_O, 0.0125 g/L FeSO_4_.7H_2_O, 0.8 g/L CaNO_3_.4H_2_O, 0.25 g/L KH_2_PO_4_ pH 6.5, 0.001% Trace Element Solution, 60 g/L mannitol, and 7 g/L agar). For each plate, 500 µl of transformed protoplasts were mixed with 500 µl of PMRT medium (PMRB medium with instead 80 g/L mannitol and 1.5 g/L agar). After 7 days, the resulting *P. patens* plants were subjected to two rounds of selection on solid complete medium supplemented with 40 µg/ml G418 to eliminate potential resistance caused by the presence of plasmid DNA not integrated into the *P. patens* genome. Mutants were identified through 2 consecutive rounds of genotyping. In the first round, primers binding the deleted region were used, and plants failing to amplify this region were selected for further analysis. For the second round, pairs of primers with one oligo binding outside the homology regions present in the vector were used to ensure the correct insertion of the plasmid into the genome. *PpATG14* expression levels in *atg14-16* and *atg14-34* lines were evaluated by RT-qPCR (see Quantification of auxin-related gene expression for details on RNA extraction, cDNA synthesis and qPCR protocol). Primers used for genotyping are listed in the supplemental table S1.

### 2.5 Determination of chlorophyll content and quantification of cell death

Total chlorophyll content was quantified in 28-day-old plants grown without cellophane on complete medium and ammonium-free medium. In this case, pigments were extracted with 80% (v/v) acetone. Chlorophylls were quantified by spectrophotometry at 664 nm and 648 nm. Determinations were performed in triplicate. Calculations were performed using the following equation: Chla ug.ml-1= [(12.25*Abs664) – (2.79*Abs648)]; Chlb ug.ml-1= [(21.5*Abs648) – (5.10*Abs664)].

For cell death quantification, 28d plants grown on solid ammonium-free medium without cellophane were incubated in 1.5-ml Eppendorf tubes with 0.05% Evans Blue for 2 h. Subsequently, four washes with distilled water were performed to remove excess stain. The dye bound to dead cells was solubilized in 50% methanol containing 1% SDS for 45 min at 50°C. Each biological replicate consisted of three plants incubated with 2 ml of the methanol/SDS mixture, and three replicates were analyzed per condition. For each replicate, a 300-µl aliquot was transferred to a well of a 96-well multiwell plate to measure absorbance at 600 nm. The dry weight of the plants was determined after completion of the procedure by drying the plants for 18 h at 65°C. Results are expressed as OD₆₀₀ per gram of dry weight.

### 2.6 Cell length measurements

Caulonemal cell length measurements were performed on subapical cells (n=20) of 7-day old filaments regenerated from protoplasts growing in ammonium-free media. Images were taken with an Olympus BX-61 microscope equipped with an Olympus DP71 color camera, and analysed using FIJI-IMAGEJ.

### 2.7 Protoplasts regeneration and caulonemata development analysis

Protoplasts from WT, *atg14-16,* and *atg5* lines were obtained following the protocol for plant transformation mentioned above. After the mannitol washes, protoplasts were resuspended in PMRT medium at a density of 2 × 10^4^ protoplasts per ml, and 1 ml of this solution was plated into Petri dishes containing PMRB medium and cellophane. After 2 days in PMRB, regenerating protoplasts were transferred to complete medium for 48 h. Next, cellophanes were moved to the following caulonemata-inducing treatments: 1) new complete medium plate (control); 2) ammonium-free medium (+NO_3_^-^, -NH_4_^+^, complete medium without di-ammonium tartrate), 3) auxin treatment (complete medium + 1 µM NAA); 4) complete nitrogen deficiency (-N: -NO_3_^-^, -NH_4_^+^, ammonium-free medium in which KNO_3_ was replaced by 1.41 g/L NaK tartrate). Tissue was observed after 2 and 4 days of treatment using an Olympus BX-61 microscope equipped with an Olympus DP71 color camera. 40 filaments were quantified per condition, and the experiment was repeated three times. For quantification, only main filaments with a minimum length of three cells were considered. Two parameters were recorded: (1) the total number of filaments containing at least one caulonemal cell (indicating caulonemal filament production), and (2) the number of caulonemal cells in each filament analyzed (indicating caulonemal filament growth).

For caulonemal filament production the effect of genotype, treatment and timepoint was modeled using a binomial generalized linear model (GLM) with the number of caulonemal filaments and non-caulonemal filaments as a two-column response variable. The full model included all main effects and their three-way interaction, and Dunnett-adjusted contrasts were used to compare each treatment against the control medium within each genotype and timepoint. Analyses were performed in R (v4.4.1) using the emmeans package.

For caulonemal filament growth, the number of filaments containing 1, 2, 3, or 4 or more caulonemal cells for each genotype is shown in a frequency distribution of total caulonemal filaments observed for each combination of treatment and time point. The distribution of the different length categories was tested by Poisson log-linear models and likelihood-ratio χ² tests.

### 2.8 IAA quantification

Hormone levels in 7d protonemal tissue from WT, *atg5*, and *atg14* lines were quantified by liquid chromatography–tandem mass spectrometry (LC–MS/MS). Tissue was grown for 4 days on complete medium with cellophane and then transferred to ammonium-free medium, -N or a new complete medium plate (control) for 3 additional days. Three biological replicates were analyzed for each genotype. For hormone extraction, 100 mg of tissue previously ground in liquid nitrogen was homogenized with 500 µl of a 1-propanol/H_2_O/concentrated HCl solution (2:1:0.002, v/v/v) by shaking at 4°C for 30 min. Subsequently, 1 ml of dichloromethane (CH_2_Cl_2_) was added to each sample, followed by further homogenization by shaking at 4°C for 30 min and centrifugation for 5 min at 13,000 × g. The lower organic phase (approximately 1 ml) was collected into glass vials and evaporated under a nitrogen stream. Samples were then re-dissolved in 0.25 ml of a solution containing 50% methanol (HPLC grade) with 0.1% formic acid and 50% H2O with 0.1% formic acid using a vortex mixer.

Chromatographic analyses were performed using an LC–MS system (Waters Xevo TQ-S Micro, Waters, Milford, MA, USA) equipped with a quaternary pump (Acquity UPLC H-Class, Waters), an autosampler (Acquity UPLC H-Class, Waters), and a reverse-phase column (Waters BEH C18, 1.7 µm, 2.1 × 50 mm). The mobile phase consisted of water with 0.1% formic acid (A) and methanol with 0.1% formic acid (B), with a flow rate of 0.25 ml/min. The initial gradient was held at 40% B for 30 s, then increased linearly to 100% B and maintained for 3 min. IAA identification and quantification were performed using a Xevo TQ-S Micro mass spectrometer (Waters, Milford, MA, USA) coupled to the UPLC system. Electrospray ionization (ESI) was used as the ion source, and mass spectra were acquired in positive ion mode. Data acquisition and processing were carried out using MassLynx software (version 4.1). The mass-to-charge (m/z) transitions monitored were 176.0 → 103.0 and 176.0 → 130.0. Quantification was performed using calibration curves with linear regression, and results are expressed as ng of phytohormone per g of fresh weight. For each condition, values for all genotypes are shown as relative to the WT in the same condition.

### 2.9 Quantification of auxin-related gene expression

Homogenized protonemata from WT, *atg5* and *atg14-16* lines were grown in complete medium for 4 days and then transferred to the following caulonemata-inducing conditions for 3 additional days: 1) new fresh complete medium plate (control); 2) complete medium + 5mM NAA (tissue was placed in a second complete medium plate for 3 days and 4.5 h prior to collection it was transferred to a third complete medium plate supplemented with NAA); ammonium-free medium (+NO_3_, -NH_4_^+^); -N (-NO_3_, -NH_4_^+^). Treatments were carried out in triplicates, and all samples were collected 7 days after homogenization. Total RNA was extracted using Quick-Zol (Kalium Technologies) according to the manufacturer’s instructions. 2 mg per sample of total RNA was treated with 2 units of DNAse I (Thermo Fisher Scientific) and used as template for cDNA synthesis. RevertAid Transcriptase (Thermo Fisher Scientific) and oligo-dTs were used according to the manufacturer’s protocol. The quantitative PCR reaction was carried out in LineGene 9600 Plus Fluorescence Quantitative Detection System using EvaGreen® (Genbiotech) and Dream Taq (Thermo Fisher Scientific). Primers used for qPCR are listed in the supplemental table S1. Actin and Ade-PRT were used as reference genes to normalize the samples. For each gene and condition, values are shown as relative to the WT in complete medium (control condition).

### 2.10 Autophagic flux assay via immunoblotting

In three independent experiments, protonema of the reporter line PpATG8epro:GFP-PpATG8e was grown on cellophane disks overlaid on solid complete medium, inside sterile 12-well multiwell plates. The tissue was grown for 4 days on complete medium (+NH_4_^+^, +NO_3_) and then transferred to solid media: complete (control), 1 μM NAA (supplemented complete), ammonium-free (-NH_4_^+^, +NO_3_^-^) and -N (-NH_4_^+^, -NO_3_^-^). Samples were harvested at the indicated time points, both at the end of the light or the dark period of the different days after transfer to treatment. Samples were ground in liquid nitrogen in 1,5 ml microtubes using pestle pellets, and 50 μl of protein extraction buffer was added (100 mM Tris-HCl, 1 mM EDTA, 2% SDS, 100 mM NaCl, 1 mM PMSF, and protease inhibitor Complete Roche). Samples were centrifuged at 13000 RPM for 10 min, the protein-containing upper phase was collected, and protein concentration was measured using the Lowry method using a SYNERGY H1 microplate reader (Biotek). Samples containing 20 ug of total protein were separated in an electrophoretic run, using 12% acrilamide SDS-PAGE gels, after which proteins were transferred to a PVDF membrane. Equal sampling was ensured by 0,5 % Ponceau S staining, and membranes were blocked for 1 hour in TBS-T buffer (50 mM Tris-HCl, 150 mM NaCl, 0,1% Tween20, pH 7.5) with 5% skimmed milk, after which membranes were washed with TBS-T once for 15 min and twice for 10 min. Membranes were incubated overnight at 4°C with anti-GFP antibody 1:5000 in TBS-T (Agrisera Rabbit αGFP AS20), washed again, incubated at RT for 1 hour with secondary antibody 1:10000 (αRabbit igG Alkaline Phosphatase Conjugate, Sigma A-3687), and subsequently washed before imaging. Lastly, membranes were incubated with an AP-revealing solution (Tris-Cl 100 mM, MgCl_2_ 5 mM, 0,033% w/v NBT, and 0,015% w/v BCIP). The membranes were imaged using the GelDoc Go Gel Imaging System (BioRad), and bands were quantified using image analysis libraries from Python (scikit-image). Autophagic flux was determined as the ratio between free GFP and the sum of free GFP and GFP-ATG8e. Flux values were normalized to T0 (D2, end of light period) of the control medium within each independent experiment. Statistical analyses were performed using linear least squares regression. Dunnett-adjusted contrasts were used to compare treatments against the control medium. Analyses were performed in R (v4.4.1) using the lme4, lmerTest, and emmeans packages.

## 3. RESULTS

### 3.1 Autophagy-deficient mutants show enhanced plant spread and sucrose accumulation under ammonium-free conditions

With the aim of evaluating caulonemata development in relation to autophagy, we first measured plant spread (an indicator of caulonemal growth) in autophagy-deficient mutants. To accomplish this, we grew equal amounts of protonemal inocula from wild type (WT) and the previously characterized *atg5* and *atg7* mutants (Pettinari et al., 2022) on medium containing nitrate as the single nitrogen source (ammonium-free medium, -NH_4+_). Although this medium does not impose severe nutritional stress for WT plants, the absence of ammonium favors the transition from chloronemata to caulonemata, thereby facilitating the observation of this developmental process. After 19 days of growth, both *atg* mutant plants exhibited increased final plant areas compared to the WT, with a significantly higher carrying capacity (K), estimated using a logistic growth model (Wald test, each mutant vs WT, p<0.001) (Figures 1-A, 2-B). This spreading response resembles the known effect of nitrogen starvation on WT plants, where caulonemata differentiation is further promoted to enhance substrate colonization and nutrient acquisition (Pettinari et al., 2022).

**Figure 1.**
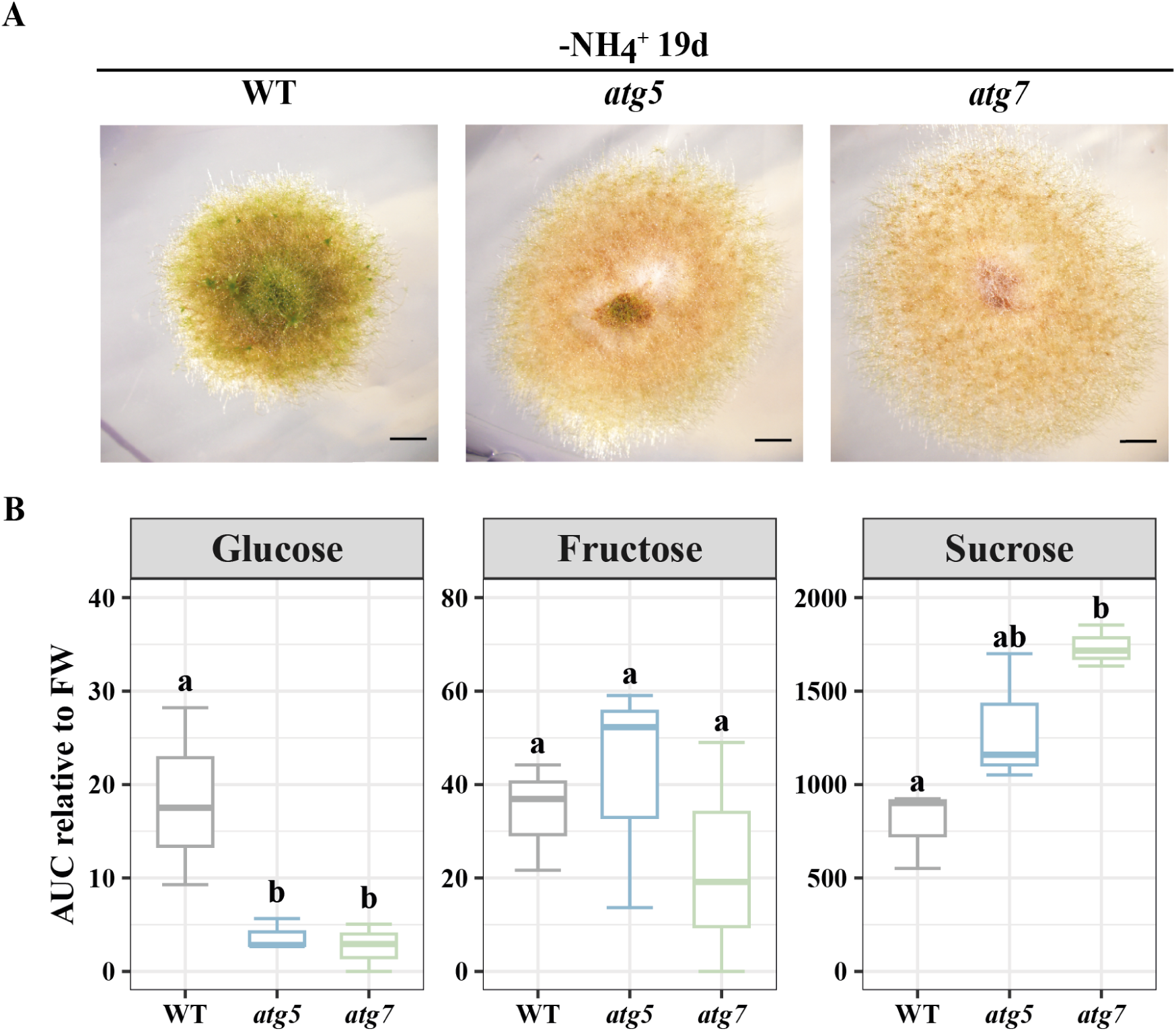
Autophagy-deficient mutants display enhanced spreading and sucrose accumulation under ammonium-free conditions. A) Plant phenotypes of WT, *atg5* and *atg7* genotypes growing in ammonium-free medium (-NH_4_^+^, +NO_3_^-^). Images were taken at the end of the experiment (19 days). Scale bar: 2 mm. B) Quantification of soluble sugars via HPLC in 7-day-old protonemata of WT, *atg5* and *atg7* lines grown in ammonium-free medium. AUC: area under the curve. Different letters indicate significant differences between genotypes (*n*= 3, One-way ANOVA and Tukey’s HSD test; p < 0.05).

**Figure 2.**
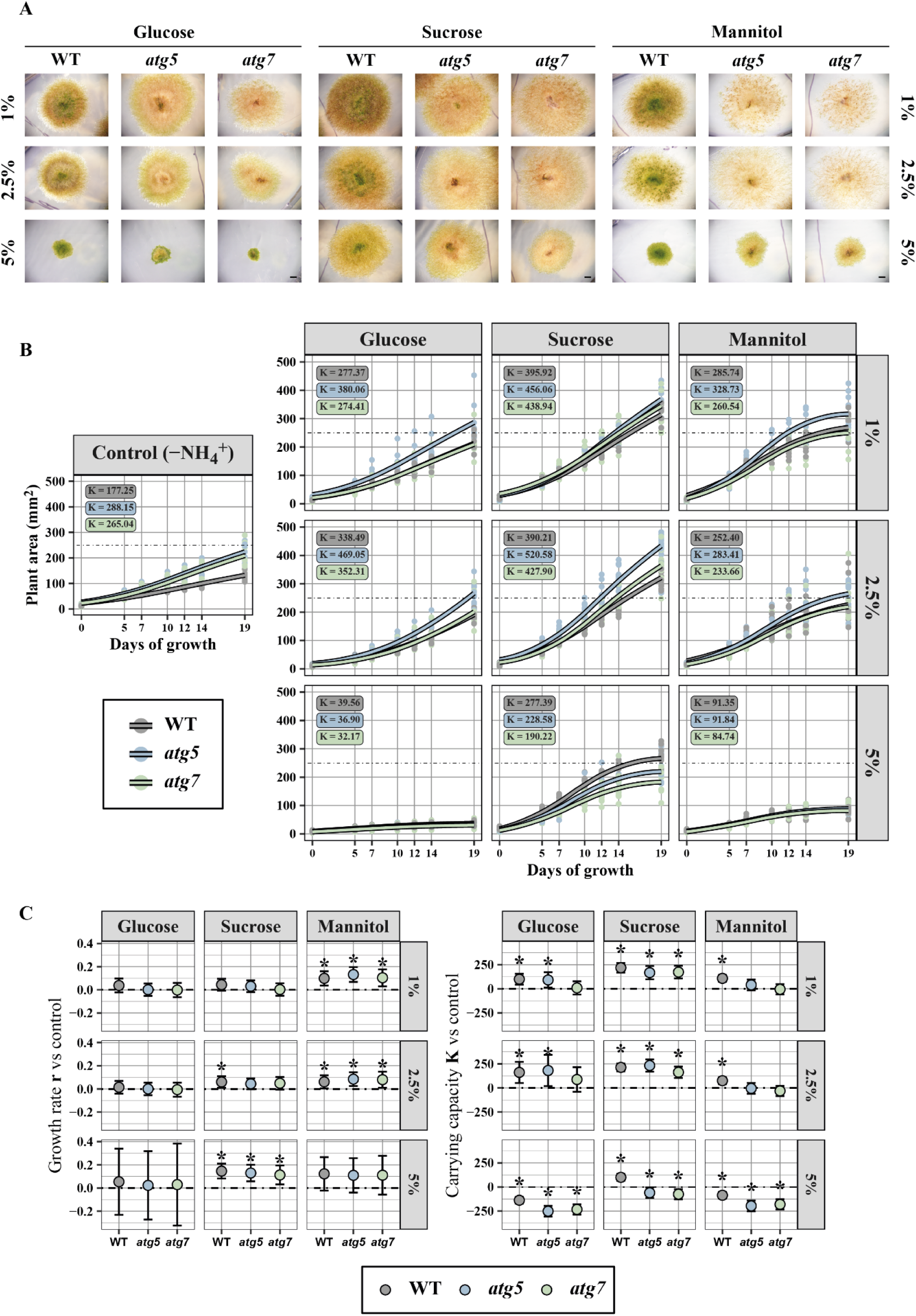
Sugar response is altered in autophagy-deficient mutants. A) Representative images of WT, *atg5* and *atg7* plants after 19 days of growth in ammonium-free medium (-NH_4_^+^, +NO_3_^-^) supplemented with 1%, 2.5% and 5% glucose, sucrose or mannitol. Scale bar: 2 mm. B) Plant growth over 19 days. Curves represent fitted values from a logistic growth model. Points represent raw plant area at each time point (n=9). Estimated intrinsic growth rate (r) values are shown within each panel. The dashed line indicates values of 250 mm2. The leftmost panel represents growth in control (non-supplemented) ammonium-free medium. C) Differences in intrinsic growth rate (r, left) and carrying capacity (K, right) between each sugar treatment and the control medium (-NH_4_^+^), within each genotype. The dashed line indicates values of 0. Error bars indicate standard error (SE). Asterisks denote significant differences in r or K between treatment and control (Wald test, p<0.05).

Auxin, sugar supplementation and nitrogen deficiency are known inducers of the chloronemata to caulonemata transition. We previously reported that *atg5* and *atg7* mutants accumulate IAA when grown on ammonium-free medium (Pettinari et al., 2022), consistent with the enhanced plant spreading response shown here. Given that autophagy deficiency leads to alterations in cellular metabolism, which includes the C/N ratio, we tested whether endogenous sugar levels are altered in these mutants. We measured glucose, sucrose, and fructose content in WT, *atg5* and *atg7* protonemata grown in ammonium-free medium. The *atg* mutants accumulated higher amounts of sucrose, whereas they exhibited lower glucose levels compared to the WT. In contrast, no significant differences were detected in fructose content between genotypes (Figure 1-B). Altogether, these results suggest that increased plant spread (and thus caulonemata development) observed in autophagy-deficient mutants under ammonium-free conditions may be partially explained by their internal sucrose accumulation, a transportable sugar known to promote caulonemata differentiation (Thelander et al., 2005).

### 3.2 The spreading response to sucrose is reduced in atg mutants

Since *atg* mutants exhibited an altered sugar profile, we tested the effect of increasing the C/N ratio through the addition of external sugars. Plant spread was evaluated in ammonium-free medium (-NH_4_^+^) supplemented with different concentrations of glucose, sucrose or mannitol (Figure 2). Plant spread was then modelled using a logistic growth model to estimate the carrying capacity (K) and the intrinsic growth rate (r) for each genotype in each medium (for details refer to section 2.2 of Materials and Methods).

Low and intermediate concentrations of glucose and sucrose significantly increased the carrying capacity (K) of most genotypes compared to the control non-supplemented medium, increasing final plant area (Figures 2-A-C). However, glucose did not significantly affect the intrinsic growth rate (r) of plants of any genotype at these concentrations, whereas higher concentrations (5% w/v) reduced K of all genotypes compared to the control, reflecting lower final plant areas (Figure 2-C). Sucrose, in contrast, significantly increased r in a concentration-dependent manner. This effect was observed only in WT at intermediate concentrations but extended to all genotypes at 5% w/v (Figure 2-C). This delayed response in the mutants was specific to sucrose and not observed with glucose. Furthermore, while the WT still displayed an increase in final plant area at 5% w/v sucrose compared to the control, the mutants instead showed a reduction, as evidenced by significant changes in K (Figure 2-C). The delay in increasing r and the reduction in K at high sucrose concentrations suggest that *atg* mutants exhibit an attenuated spreading response that is specific to sucrose compared to the WT.

Mannitol was included as an osmotic control. At low and intermediate concentrations all genotypes grew faster (r) but only the WT reached higher final areas (higher K) compared to the control medium (Figures 2-B, 2-C). This response differed from that observed with glucose or sucrose, indicating that these sugars have specific effects not solely attributable to an osmotic effect. At higher mannitol concentrations all genotypes exhibited reduced plant areas, suggesting that the response to 5% w/v glucose is largely osmotic. It is relevant to note that, at the same sugar percentages, glucose and mannitol exert almost double the osmotic effect compared to sucrose (Figure 2).

The increased spread and sucrose accumulation of the *atg* mutants in the non-supplemented ammonium-free medium (-NH_4_^+^, Figures 1, 2) suggest that, under these conditions, the mutants exhibit an elevated basal spreading state associated with a high C/N ratio. They show a reduced spreading response to sucrose, but not glucose (Figure 2), and accumulate sucrose, but not glucose (Figure 1-B). Together, these observations suggest that exogenous sucrose does not elicit an additional response in the mutants because their spreading state is already elevated in the non-supplemented medium. Altogether, these results suggest that autophagy contributes to the coordination of plant spread (mainly regulating caulonema development) in response to changes in carbon and nitrogen availability (C/N ratio), thereby impacting the internal sugar balance.

### 3.3 Autophagy-deficient mutants show increased, but ultimately unsustainable, caulonemata growth under nitrogen-deficient conditions

Since *PpATG5* and *PpATG7* encode proteins that participate in the same step of the autophagic process (i.e. ATG8 lipidation), to strengthen our analysis, we chose an additional *ATG* gene involved in a different step of autophagosome biogenesis. We generated an independent knockout line by disrupting PpATG14, which encodes the differential subunit of the PI3K complex I involved in the decoration of the autophagosome membrane with PI3P (Gross et al., 2025). Mutants lacking PpATG14 showed an intermediate senescent phenotype between the WT and the previously characterized *atg5* genotype (Pettinari et al., 2022). This was observed both in young plants regenerated from protoplasts after 7 days of growth in complete media (Figure 3-A) as well as in mature plants grown under both complete and ammonium-free (-NH_4_^+^) media (Figure 3-B). Quantification of chlorophyll content (Figure 3-C) and cell death (Figure 3-D) showed that *atg14* plants exhibit a less severe, yet consistent, premature senescence compared to *atg5*. As we previously reported for *atg5* (Pettinari et al., 2022), caulonemal cell length was significantly reduced in *atg14* plants compared to the WT (Figure 3-E). Out of the two independent *atg14* lines (*atg14-16* and *atg14-34*), *atg14-16* showed the most drastic reduction in *ATG14* transcript levels (Figure S1). Therefore, *atg14-16* mutants were included together with the *atg5* line in subsequent experiments.

**Figure 3.**
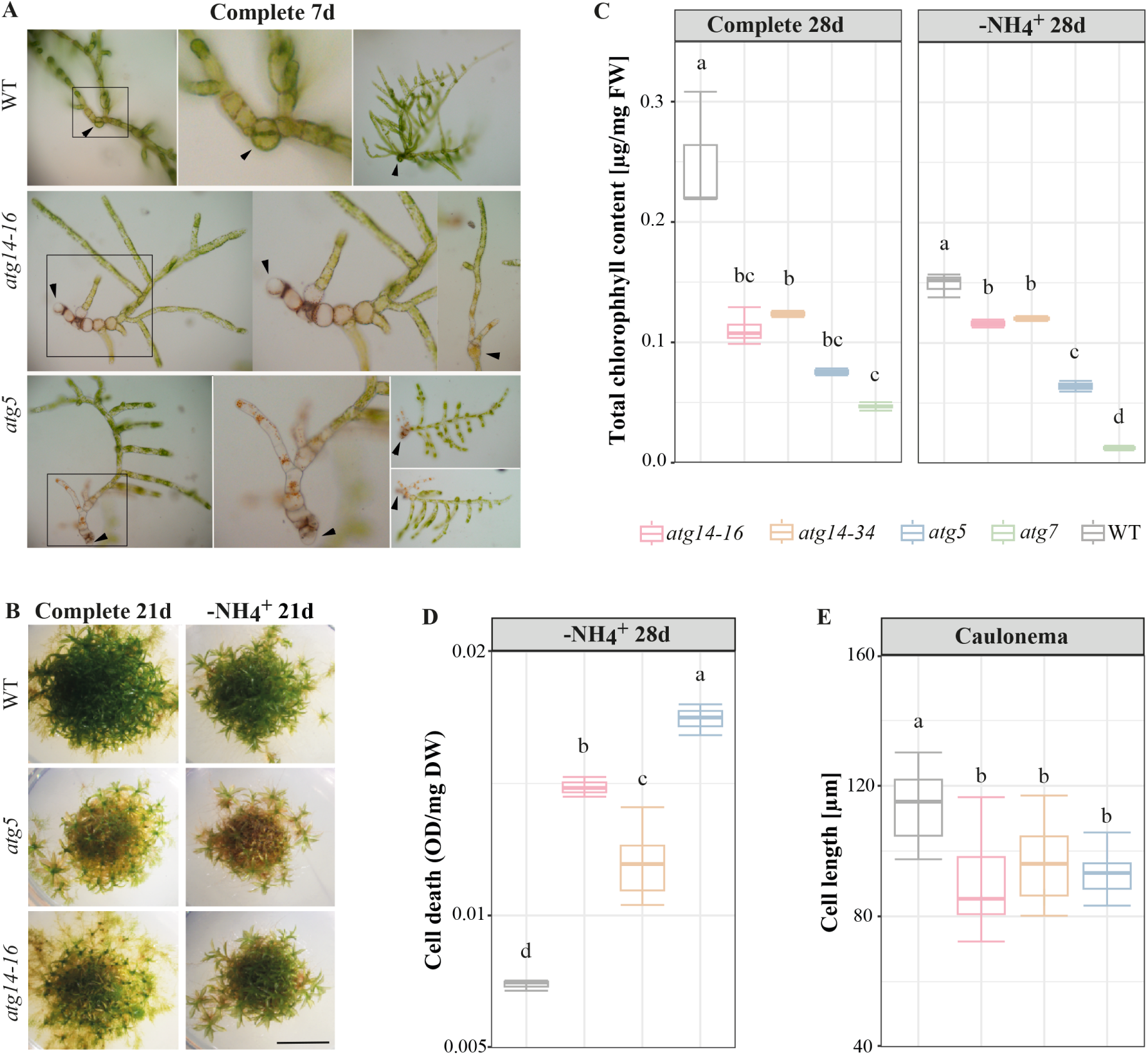
Mutants lacking PpATG14 show an intermediate phenotype between the WT and *atg5* mutants. A) Representative images of filaments regenerated from protoplasts after 7 days of growth in complete medium. Squares highlight senescent areas in the center part of the regenerating filaments from *atg5* and *atg14* genotypes, in contrast to the WT. Black arrowheads indicate the oldest cell in the filament. B) Representative images of WT, *atg5* and *atg14* plants after 21 days of growth in complete (+NH_4_^+^, +NO_3_^-^) and ammonium-free (-NH_4_^+^, +NO_3_^-^) media. Scale bar: 6 mm. C) Total chlorophyll content in WT, *atg5*, *atg7* and two independent *atg14* (*atg14-16* and *atg14-34*) lines. Quantification was carried out in colonies after 28 days of growth in complete and ammonium-free media. Different letters indicate significant differences between genotypes (*n* = 3, One-way ANOVA and Tukey’s HSD test; P < 0.05). D) Cell death quantification using Evans Blue in WT, *atg5* and *atg14* colonies after 28 days of growth in ammonium-free medium. Different letters indicate significant differences between genotypes (*n* = 3, One-way ANOVA and Tukey’s HSD test; P < 0.05). E) Cell length measurements of subapical caulonemata cells growing in ammonium-free conditions. Different letters indicate significant differences between genotypes (*n*= 20, One-way ANOVA and Tukey’s HSD test; P < 0.05).

To accurately assess caulonemata development in *atg* mutants, we regenerated plants from protoplasts. By initiating growth from a single cell, plants are developmentally staged and it is possible to more readily observe the chloronemata-to-caulonemata transition. In this case, we tested the effects of increasing the C/N ratio by lowering nitrogen availability in the media instead of adding sugars. Protoplasts were regenerated in a complete medium supplemented with 6% mannitol for 5 days to allow cell wall regeneration and initial chloronemal growth. Then, regenerated plants were transferred to two different caulonemata-inducing conditions: ammonium-free medium (-NH_4_^+^), and complete nitrogen deficiency (-N). Transferring to fresh complete media was used as a control. To account for caulonemal filament production, the percentage of primary filaments showing caulonemal identity was quantified 2 and 4 days after transferring (Figure 4-A). Additionally, to assess progression of caulonemal filament growth, we classified caulonemal filaments in 4 categories, ranging from 1 to 4+ caulonemal cells, and quantified the number of caulonemal cells per filament (Figure 4-B). The earlier the chloronema-to-caulonema transition occurs, the higher number of caulonemal cells per filament is observed, enabling identification of genotypes or conditions that trigger this transition faster.

**Figure 4.**
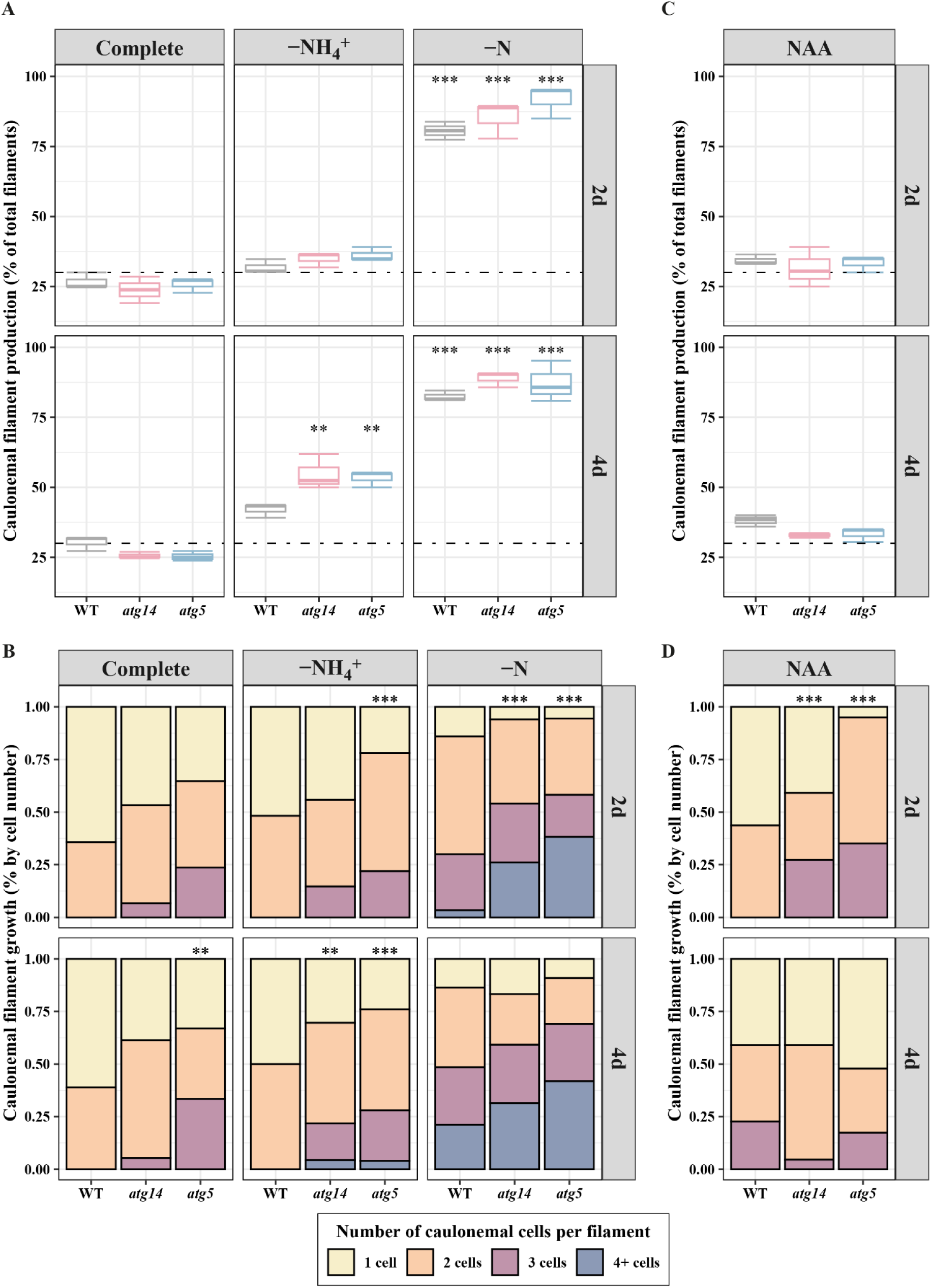
Caulonemal filament production and growth in *atg* mutants is modulated differently upon nitrogen deficiency and auxin treatment. A) Caulonemal filament production expressed as percentage of total filaments in WT, *atg14* and *atg5* after 2 or 4 days of transferring the regenerating protoplasts to complete (+NH_4_^+^, +NO_3_^-^, control), ammonium-free (-NH_4_^+^, +NO_3_^-^) and -N (-NH_4_^+^, -NO_3_) media. Asterisks denote significant differences between treatment and control (complete) media within each genotype and timepoint, as determined by Dunnett contrasts from a binomial GLM (*** p < 0.001, ** p < 0.01). B) Caulonemal filament growth, expressed as the frequency distribution of caulonemal filaments with a given number of caulonemal cells after transfer to complete, ammonium-free or -N media. Caulonemal filaments identified in A were categorized according to the number of caulonemata cells present in each filament. Asterisks indicate a significant effect of the genotype on the distribution of the different length categories (1 cell, 2 cells, 3 cells, 4 or more cells) as tested by Poisson log-linear models and likelihood-ratio χ² tests (*=P < 0.1, **= P < 0.05, ***= P < 0.01). C) Caulonemal filament production expressed as percentage of total filaments in WT, *atg14* and *atg5* after 2 or 4 days of transferring the regenerating protoplasts to complete medium supplemented with 1μM NAA. Statistical analysis was performed as in A. D) Caulonemal filament growth, expressed as the frequency distribution of caulonemal filaments with a given number of caulonemal cells after transfer to complete medium supplemented with 1μM NAA. Statistical analysis was performed as in B.

In the complete medium, no significant differences in caulonemal filament production were observed between genotypes at any analyzed timepoint (Figure 4-A, complete). However, filaments with up to 3 caulonemal cells were found in the *atg* mutants in the complete medium while WT plants only showed filaments with 1 and 2 caulonemal cells (Figure 4-B, complete medium). This observation suggests that, in the complete medium, *atg* mutants initiate the transition from chloronema to caulonema earlier, although this does not translate into a higher caulonemal filament production. Earlier differentiation in the complete medium was also observed in 3D structures of *atg* mutants, such as buds and gametophores (Figure 5-A, left panel), indicating that autophagy-deficient genotypes exhibit an overall accelerated life cycle progression and, consequently, premature senescence (Figure 5-A, right panel, Pettinari et al., 2022).

**Figure 5.**
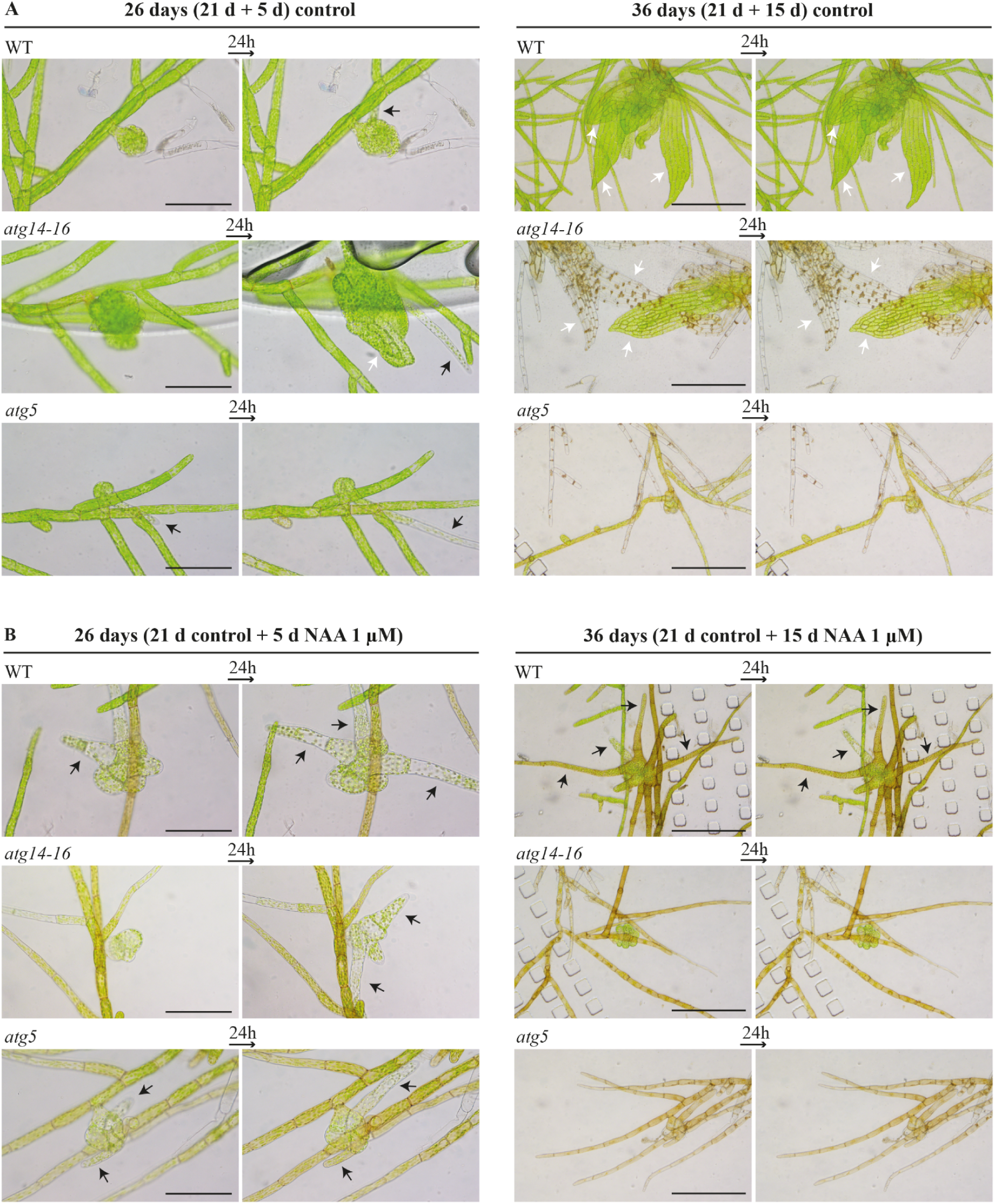
*atg* mutants display hypersensitivity to auxins. Tissue from WT, *atg14-16*, and *atg5* lines grown in microfluidic chambers for 21 days under control conditions and for an additional 5 (left panels) or 15 (right panels) days on Hoagland liquid medium (A) or Hoagland supplemented with 1 μM NAA (B). Scale bars: 200 μm (left panels) and 500 μm (right panels). Black arrows indicate rhizoids, and white arrows indicate phyllids.

After 2 days of growth in the ammonium-free condition, no significant differences in caulonemal filament production were detected compared to the complete medium for any genotype (Figure 4-A, -NH_4_^+^). Likewise, no differences were detected regarding caulonemal filament growth compared to the control medium, although the *atg* mutants, as in the complete medium, exhibited higher growth compared to the WT (Figure 4-B). On the other hand, after 4 days of growth in the ammonium-free condition both *atg5* and *atg14* exhibited significantly higher caulonemal filament production compared to the control medium, while the WT did not (Figure 4-A). Furthermore, after 4 days, both mutants showed enhanced caulonemal filament growth (some with 4 or more cells) than the WT, an effect being even more pronounced than in the control medium (Figure 4-B).

Complete nitrogen deficiency (Figure 4-A, -N) induced a significant increase in the production of caulonemal filaments of all genotypes after just 2 days of growth and similar values were observed after 4 days. These observations suggest that differential caulonemal production occurs mostly in the first 2 days of exposure to complete nitrogen deficiency. Noteworthily, after 2 days, caulonemal filament growth in the *atg* lines was significantly higher than the WT (Figure 4-B, -N), indicating that the caulonemal-inducing response to nitrogen deficiency is amplified in the absence of autophagy, at least in the short term. Although the 4 categories of caulonemal filaments were present in all genotypes, the distribution was significantly different, with *atg* plants showing a higher proportion of filaments with 4 or more caulonemal cells (Figure 4-B, -N 2d). Interestingly, after 4 days, while the number of filaments with 4 or more caulonemal cells increased in the WT, no such growth was observed in the mutants, thus erasing the initial genotypic differences. This growth stagnation in the mutants suggests that, although caulonemata development is rapidly induced under complete nitrogen deficiency, further elongation of the existing filaments appears unsustainable over time.

### 3.4 atg mutants are unable to sustain growth under prolonged auxin exposure

Caulonemal development is tightly regulated by auxin signaling. To explore this modulation in the context of autophagic deficiency, we regenerated plants from protoplasts and evaluated caulonemal development after transferring them to complete medium supplemented with 1 μM NAA (1-naphthaleneacetic acid). As in the previous experiment, we quantified caulonemal filament production (Figure 4-C) and caulonemal filament growth (Figure 4-D).

No significant changes in the production of caulonemal filaments were observed either after 2 or 4 days of growth in the presence of NAA (Figure 4-C, NAA). After 2 days, caulonemal filament growth was similar for all genotypes compared to the complete medium, although auxin supplementation reinforced the already observed difference between WT and *atg* mutants (Figure 4-D, NAA 2d). Intriguingly, after 4 days, differences in caulonemal filament growth between genotypes were no longer detected (Figure 4-D, NAA 4d). This convergence could be a consequence of an apparent stagnation of caulonemal growth (between 2 and 4 days) in the mutants compared to the WT and suggests that further growth may be unsustainable. The *atg* mutants phenotype resembles that observed in response to nitrogen deficiency (Figure 4B). However, unlike nitrogen deficiency, NAA treatment did not enhance the production of caulonemal filaments in the mutants (Figure 4-D). These observations are further supported by an independent experiment showing that WT and *atg* mutant plants growing in microfluidic devices responded similarly to short-term auxin treatment, while prolonged exposures led to growth arrest and the presence of dead cells in the mutants (Figure 5-B).

### 3.5 Endogenous auxin homeostasis is altered in atg mutants

The altered response to NAA supplementation in the *atg* genotypes over time suggests that endogenous auxin homeostasis is disrupted in mutants with impaired autophagic activity. This is consistent with our previous work showing IAA accumulation in *atg5* and *atg7* plants grown in ammonium-free medium for 7 days (Pettinari et al., 2022). To assess this in the context of variable nitrogen availability, we quantified IAA levels in *atg5* and *atg14* protonema grown on complete medium for 4 days and then transferred to ammonium-free medium (-NH_4_^+^) or complete nitrogen deficiency (-N) for 3 additional days. Transferring to fresh complete media was used as a control. In complete medium, all genotypes showed similar IAA levels (Figure 6). However, after transferring to both nitrogen-limiting media, a significant accumulation of IAA was detected in the *atg* mutant lines compared to the WT (Figure 6, -NH_4_^+^ and -N), with the highest values observed in complete nitrogen deficiency. Notably, *atg5* consistently showed the highest IAA levels, while *atg14* exhibited intermediate values in both conditions, which is consistent with *atg14*’s intermediate phenotype (Figure 3). These results point to alterations in endogenous auxin homeostasis being linked to changes in the C/N supply ratio in the absence of autophagy.

**Figure 6.**
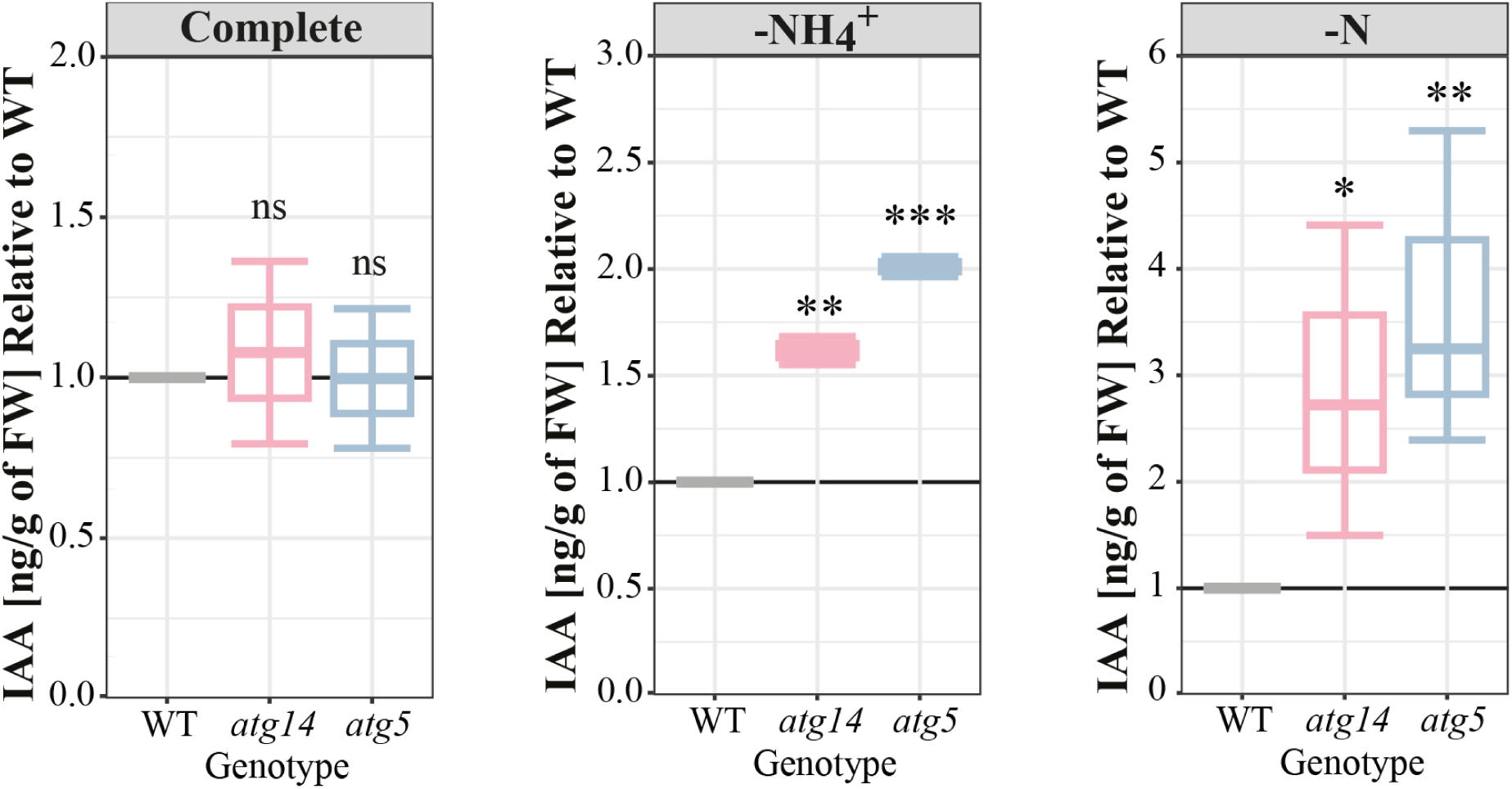
IAA accumulates in *atg* mutants after transfer to mild and complete nitrogen deficiencies. IAA content in protonemata was measured after 3 days of transfer to complete (+NH_4_^+^, +NO_3_^-^, control), ammonium-free (-NH_4_^+^, +NO_3_^-^) and -N (-NH_4_^+^, -NO_3_) media. For each condition, asterisks indicate significant differences between each *atg* mutant and the WT (*n*= 3, T test; *= P < 0.1, **= P < 0.05, ***= P < 0.01). In each treatment, values for every genotype are shown as relative to the WT within the same treatment.

### 3.6 Auxin-related gene expression is altered in autophagy-deficient mutants

To elucidate the cause behind the observed changes in IAA levels and caulonemata production in *atg* mutants, we evaluated the expression of auxin-related genes under both, complete medium and after transferring to ammonium-free medium (-NH_4_^+^) (Figure 7). As an additional control, tissue was incubated with 5 μM NAA for 4.5 h to verify transcriptional responses to exogenous auxin (Bascom et al., 2023). Gene expression analysis was also attempted after transferring the tissue to complete nitrogen deficiency. However, in this condition, reference genes (*PpACT* and *PpADE-PRT*) showed markedly altered Ct values in the *atg* mutants, preventing reliable normalization and quantitative interpretation of the data. Given that autophagy-deficient mutants senesce rapidly under this condition, these changes likely reflect widespread transcriptional disruption rather than specific regulatory effects. Therefore, data from this condition were excluded from further analysis. We selected markers for auxin biosynthesis (*PpSHI1*, *PpYUCb*), transport (*PpPINA*), response (*PpGH3-1*) and auxin-induced caulonemata development (*PpRSL1* and *PpRSL2*, members of the class I *RSL*, and *PpRSL4* representing class II *RSL*).

**Figure 7.**
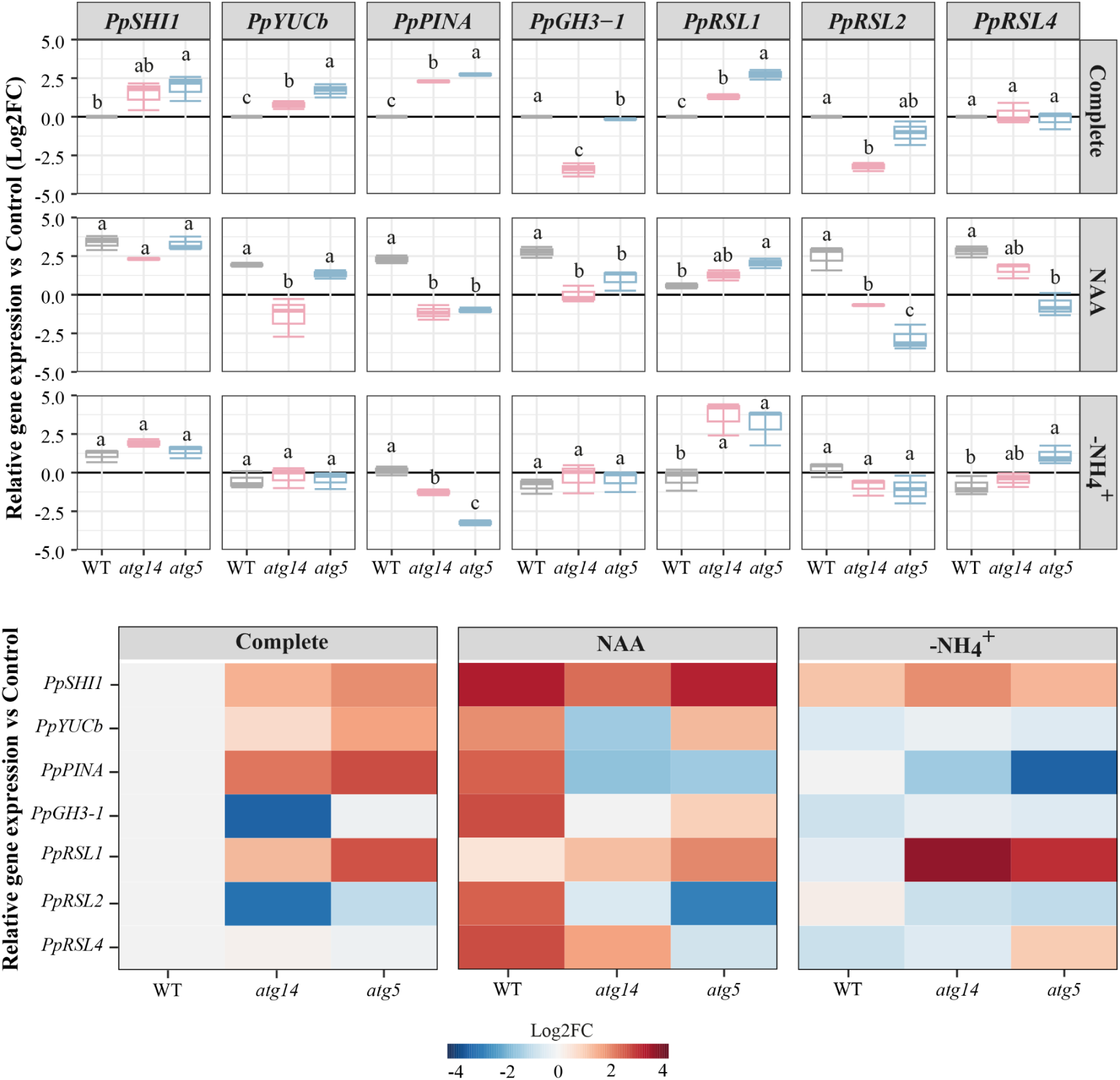
Expression of auxin-related genes is altered in mutants with impaired autophagy. Quantification of gene expression by RT-qPCR in WT, *atg5* and *atg14* protonemata after transference to complete medium (+NH_4_^+^, +NO_3_^-^), complete medium supplemented with NAA 5μM, or ammonium-free (-NH_4_^+^, +NO_3_^-^) medium. For each gene and condition, Log2FC was calculated as relative to the WT in complete medium. Upper panel: box plot showing the results of the statistical analysis; for each gene, different letters indicate significant differences between genotypes within each condition (*n*= 3, One-way ANOVA and Tukey’s HSD test; P < 0.05). Lower panel: heatmaps showing changes in relative gene expression (mean Log2FC values).

In the complete medium (Figure 7, complete), expression of the auxin biosynthesis-related genes *PpSHI1* and *PpYUCb* was higher in *atg* mutants compared to the WT, with *atg5* showing the strongest induction. Expression of the auxin transporter *PpPINA* was also increased in *atg14* and further elevated in *atg5* relative to the WT. In contrast, the auxin-responsive gene *PpGH3-1* was strongly downregulated in *atg14* and slightly reduced in *atg5* compared to the WT. Analysis of *RSL* genes revealed divergent responses among class I members: *PpRSL1* expression was increased in both *atg* mutants, with higher levels for *atg5*, whereas *PpRSL2* expression was reduced, particularly in *atg14*. Expression of the class II gene *PpRSL4* did not differ significantly between genotypes. The simultaneous increase in auxin biosynthesis-related genes and the auxin exporter *PpPINA* suggests a compensatory mechanism that maintains stable IAA levels in *atg* mutants. These results are consistent with the fact that no differences in IAA levels were found among genotypes in complete medium (Figure 5).

Following treatment with exogenous auxin (Figure 7, NAA), WT plants displayed strong induction of all analyzed genes, confirming a robust transcriptional response to auxin. In contrast, the response profile was altered in *atg* mutants. While *PpSHI1* was comparably induced in all genotypes, *PpYUCb* increased similarly to the WT in *atg5* but was reduced in *atg14*. Notably, *PpPINA* expression was significantly reduced in both *atg* mutants. *PpGH3-1* also showed lower expression in the mutant lines. Regarding *RSL* genes, NAA application increased *PpRSL1* expression in both *atg* mutants but significantly decreased *PpRSL2* levels. *PpRSL4* expression was also reduced in the mutants relative to the WT, with *atg5* showing even slightly lower levels than the control condition. Interestingly, for all *PpRSL* genes, *atg14* exhibited intermediate expression between the WT and *atg5*. These observations indicate an impaired transcriptional response to exogenous auxin in the autophagy-deficient mutants, particularly affecting auxin transport and response genes.

After three days in the ammonium-free medium (Figure 7, -NH_4_^+^), *PpSHI1* expression increased similarly in all genotypes, whereas *PpYUCb* expression remained close to levels in the complete medium, with no significant differences between genotypes. *PpPINA* expression was essentially unchanged in the WT but significantly reduced in both *atg* mutants, with the strongest decrease observed in *atg5*. *PpGH3-1* levels did not differ between genotypes. Eliminating ammonium from the media did not alter expression of class I *RSL* genes in the WT and only subtly decreased *PpRSL4* levels. However, an interesting response was observed in the mutants: *PpRSL1* expression strongly increased in both *atg* lines, whereas *PpRSL2* levels remained similar to the WT. *PpRLS4* expression was increased in *atg5* relative to the WT, and intermediate levels were found in *atg14*. Although PpPINA protein activity or accumulation was not directly analyzed, the reduction of *PpPINA* transcript levels in *atg* mutants likely contributes to the IAA accumulation previously observed under ammonium-free conditions (Figure 5).

Altogether, these results show that in the absence of autophagy, auxin homeostasis and signaling are differentially affected when plants are exposed to distinct caulonemata inducing conditions, which correlate with the observed differences in caulonemata production between WT and autophagy-deficient lines.

### 3.7 Autophagic activity diminishes in caulonemata-inducing conditions

As shown by the results presented above, caulonemata development induced by nitrogen deficiency (either by the removal of ammonium or by both ammonium and nitrate) is strongly enhanced in *atg* mutants with impaired autophagic activity. However, their response to exogenous auxin is impaired compared to the WT. To understand the effect of these caulonemata inducing stimuli on autophagic activity, we quantified autophagic flux under such conditions using a GFP-ATG8 cleavage assay in the PpATG8e::GFP-PpATG8e reporter line (Sanchez-Vera et al., 2017; Pettinari et al., 2022). This line expresses an N-terminal GFP tag fused to PpATG8e, under its native promoter. As autophagosomes are transported to the vacuole and their cargo is degraded, GFP is cleaved from ATG8e and accumulates due to its relative resistance to vacuolar degradation. Using anti-GFP antibodies, it is possible to quantify (via immunoblot) both the GFP–ATG8e fusion protein and free GFP, and autophagic flux is then estimated from the ratio between free GFP and total GFP, a widely used method to monitor autophagy (Pettinari et al., 2022; Robert et al., 2021, 2025).

For this assay, protonemal tissue was grown for 3 days in complete medium and then transferred to different caulonemata inducing media: control (complete medium), auxin supplemented medium (complete + 1 μM NAA), ammonium-free medium (-NH_4_^+^) or complete nitrogen deficiency (-N). Samples were collected across the second to fourth day after transferring, at the end of both the light and dark periods of each day (long-day photoperiod, 16:8h) (Figure 8). In the complete medium, as expected, autophagic flux was induced after each dark period, when the ratio of C/N supply was lowest (low C, high N). Exogenous auxin supply (NAA) led to a significant global reduction of autophagic flux compared to the control complete medium (p<0.01, Dunnet test, using a linear model with contrasts averaged across time), an effect most clearly visible at the end of each dark period. On the other hand, the effect of ammonium deficiency (-NH_4_^+^) had no clear pattern, and no significant effect was observed compared to the complete medium. Finally, following transfer to complete nitrogen deficiency (-N), autophagic flux was also globally reduced compared to the complete medium (p<0.01), this effect being strongest as well by the end of each dark period. In this medium (low N), during the light period (high C), the C/N supply ratio is increased, correlating with an induction of autophagic flux. Conversely, after the dark period (low C), the C/N ratio is restored to a baseline level, and autophagy flux is reduced. These results suggest that alterations to the C/N ratio induce autophagic flux, beyond the individual levels of C and N supply by themselves. Altogether, these observations indicate that autophagy is more active across the day-night cycle in nitrogen-rich conditions, while the strongest caulonemal growth-inducing conditions tested (NAA and -N, as seen in the WT in Figure 4) reduce autophagic activity.

**Figure 8.**
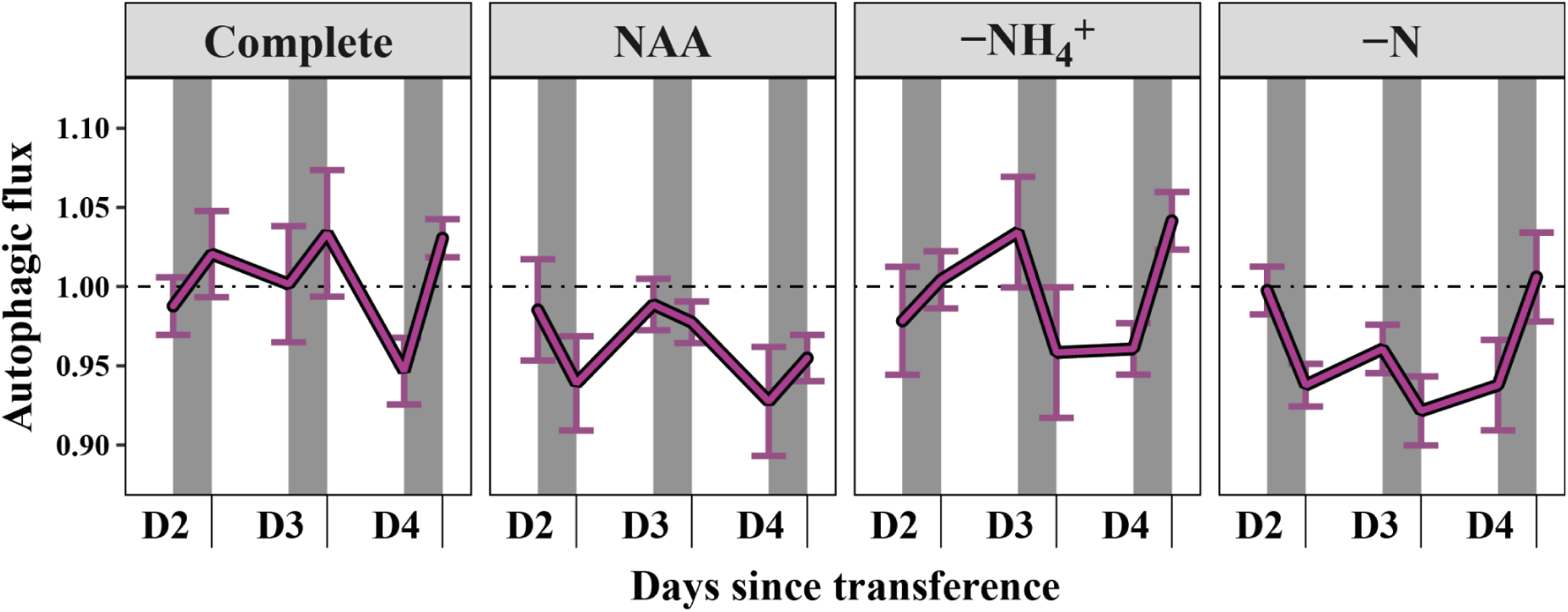
Autophagic flux decreases after transfer to caulonemata-inducing conditions. Autophagic flux modulation after transfer to complete medium (+NH_4_^+^, +NO_3_^-^, control), 1 μM NAA (supplemented control), ammonium-free (-NH_4_^+^, +NO_3_^-^) and -N (-NH_4_^+^, -NO_3_^-^), measured using a GFP-ATG8e cleavage assay. Each line represents mean autophagic flux across time, normalized to T0 (D2, end of light period) of each control within each independent experiment. Error bars indicate SE. Light and dark periods of a 16/8h day/night cycle are represented by the white and grey areas, respectively.

## 4. DISCUSSION

The integration of nutrient status with developmental programs is essential for plant survival in fluctuating environments. Here we show how the chloronema-to-caulonema transition induced by nitrogen deficiency (increasing the C/N supply ratio) is enhanced in autophagy-deficient mutants, highlighting an increased response of these genotypes to N availability, even if caulonemal growth appears ultimately unsustainable. The absence of ammonium by itself was sufficient to trigger a pronounced developmental change in these mutants. In contrast, mutants growing in ammonium-free conditions supplemented with sucrose (also increasing the C/N supply ratio) showed a lower spreading response compared to the WT. These different responses to a high C/N supply ratio might be explained because of intrinsic disruptions in the metabolism of the mutants, including high levels of internal sucrose and lower glucose compared to the WT (i.e. a basally high C/N state and thus spread). In addition, previous studies showed impaired nitrogen utilization in *P. patens* autophagy-deficient *atg3* mutants (Chen et al., 2020). These results point to nitrogen limitation heavily disrupting carbon metabolism in autophagic-deficient mutants, which probably contributes to strengthening their basal spreading through caulonemal development and diminishing their plasticity to a changing C/N supply.

Upon nitrogen deficiency, WT *P. patens* plants can deploy different strategies to obtain nutrients. On one hand, they might activate nutrient remobilization from source tissues (e.g., leaflets and chloronemal cells) to sink tissues (e.g., caulonemal cells and rhizoids) to sustain their growth. On the other hand, they could also alter their developmental program and promote caulonemata formation through the modulation of auxin signaling, thereby increasing substrate exploration. In either case, this likely involves the activation of degradation routes of high nitrogen containing organelles, like chlorophagy. Although autophagy-independent routes for chloroplast degradation have been reported in vascular plants (Wang & Blumwald, 2014; Zhuang & Jiang, 2019), there is no evidence yet that this is the case for *P. patens*. Consequently, if the plant facing this condition is an autophagy-deficient mutant, nutrient remobilization is most likely severely compromised. To compensate, *atg* plants cope with nutrient deficiency by increasing auxin-induced caulonemata development, as evidenced in protoplast-regenerated plants. Since the photosynthetically active chloronemata have abundant chloroplasts, structures rich in nitrogen which are more prone to redox damage under adverse conditions, increasing caulonemata development when facing a nitrogen deficiency probably serves as a strategy for *atg* mutants not only to search for nutrients but also to conserve nitrogen and mitigate oxidative damage. Yet, this compensatory strategy most likely represents a short-term trade-off and would prove ultimately unsustainable, as the energetic and metabolic imbalances caused by impaired autophagic activity inevitably lead to premature senescence and cell death. This energetic imbalance is also evident in *P. patens* double mutants of the energy sensor SnRK1 (*Ppsnf1a-Ppsnafb*), which activates autophagy and inhibits the TOR complex (Thelander et al., 2004, Soto-Burgos & Bassham, 2017). *Ppsnf1a-Ppsnafb* mutants display several developmental abnormalities, such as smaller gametophores, early senescence, shorter chloronemal cells, increased caulonemata development in ammonium-free medium, reduced cytokinin sensitivity, enhanced auxin sensitivity (Thelander et al., 2004). This phenotype closely resembles that of the *atg* mutants, although it is more severe in the *SnRK1*-deficient lines (Pettinari et al., 2022; Mukae et al., 2015; this work). Additionally, *tps*-deficient mutants which show lower caulonemal development and reduced sensitivity to exogenous sucrose and NAA (Phan et al., 2020), exhibit an opposite basal phenotype to the *atg* mutants. This is consistent with the fact that TPS produces T6P, a sugar-sensing metabolite and inhibitor of SnRK1, and thus a negative regulator autophagy. Their reduced response to sucrose and NAA, similarly to *atg* mutants, further contributes to the notion that sugar balance is key in regulating the chloronema-to-caulonema transition.

*atg* mutant plants regenerated from protoplasts and transferred to auxin-supplemented medium failed to sustain caulonemata development over time, and prolonged hormone exposure led to a generalized negative effect on growth. This outcome is similar to what was observed under complete nitrogen deficiency (Pettinari et al., 2022; this work). In this context, measuring auxin levels and analyzing auxin-related gene expression in different nutritional conditions provided insights into how hormonal cues shape developmental responses to nutrient availability in *atg* mutants. Results showed that these mutants accumulate IAA in the ammonium-free condition, with further increases observed under complete nitrogen deficiency. Although gene expression could not be assessed under full nitrogen starvation, transfer to the ammonium-free condition led to reduced *PpPINA* expression in *atg* mutants. If this reduction translates into decreased PpPINA activity, it could limit auxin export and result in intracellular auxin accumulation, thereby explaining the elevated IAA levels detected in these genotypes. Moreover, in the ammonium-free medium, *PpGH3-1* expression was similar between genotypes whereas *PpRSL1* was strongly upregulated in the mutants. *PpRSL4* was also induced in *atg5*, with intermediate levels in *atg14*. These findings suggest that, in autophagy-deficient plants growing under ammonium-free conditions, auxin-responsive gene regulation is conserved while caulonemata development is enhanced. Although contributions from additional class II RSL genes cannot be excluded, this developmental outcome is likely driven by a decrease in auxin export leading to IAA accumulation and subsequent induction of *PpRSL1*, with a secondary role from *PpRSL4*. Interestingly, under nutrient-rich conditions, a different regulatory pattern emerges: the reduction in *PpGH3-1* expression indicates altered auxin responsiveness, and the differences observed between both *atg* mutants suggest specificity in how distinct components of the autophagic machinery influence auxin signaling. Moreover, the opposite regulation of class I RSL genes (*PpRSL1* and *PpRSL2*) in the complete medium points to the existence of buffering mechanisms that help maintain controlled caulonemata development when nutrients are not limiting, consistent with results obtained in the plant spreading analysis and protoplast regeneration.

As in the ammonium-free condition, treatment with exogenous auxin also caused a reduction in *PpPINA* expression in the mutant lines compared to the WT. However, the ammonium-free condition enhanced the mutant caulonemal filament production and growth to a greater extent than in the WT, while treatment with auxin did not cause such differences between genotypes. This apparent incongruity could be explained by the observed differences in the expression of the auxin-responsive *PpRSLs* genes. Additionally, it is well known that the effect of exogenous auxin supply on plant growth depends on concentration, duration and stage at which the treatment is applied (Asghar et al., 2023; Thelander et al., 2018). In *atg* plants, reduced PIN-mediated auxin efflux likely leads to increased intracellular auxin accumulation, a condition probably further enhanced by NAA supplementation. Together, these effects may elevate intracellular auxin levels beyond the growth-promoting threshold, which, coupled with typical higher proximal auxin sensing observed in cells of older protonemata filaments (Thelander et al., 2019), could account for the observed negative effect on cell survival observed under prolonged exposures. In fact, it has been reported that overexpression of *PpSHI1* leads to an early senescence phenotype resembling that of *atg* mutants (Eklund et al., 2010). Considering that *atg* mutants growing in complete nitrogen deprivation showed the highest levels of IAA, auxin accumulation could also be contributing to the accelerated cell death of autophagy-deficient lines in this condition. Alternatively, auxin production could increase in dying cells as a consequence of tryptophan release during programmed cell death (PCD) (Sheldrake, 2021). Although autophagy is the main mechanism for PCD in plants, it has been reported that the 26S proteasome pathway is enhanced in *P. patens* mutants lacking *ATG3* to compensate for autophagy deficiency (Chen et al., 2020). Thus, ubiquitin-mediated protein degradation could be a possible source of tryptophan release and auxin production in the absence of autophagy. Whether IAA accumulation is a cause or a consequence of early senescence in *atg* mutants remains unexplored.

Of note, early senescence and developmental alterations in *atg14* were less severe than in *atg5*. This is puzzling, as phagophore decoration with PI3P (mediated by ATG14 as the autophagic subunit of PI3K complex I) and ATG8 lipidation (involving ATG5) are both key events during autophagosome biogenesis (Gross et al., 2025). However, this is not the first time that differences in the severity of the senescent phenotype have been observed among mutants of the autophagic machinery in other plants, which does not necessarily correlate with the stage of the process in which each protein participates (Kang et al., 2018; Li et al., 2014; Liu F. et al., 2020; Suttangkakul et al., 2011). Interestingly, the phenotypic differences between *atg5* and *atg14* correlate with the auxin-related results observed in these mutant lines. Under the ammonium-free condition, IAA levels in *atg14* mutants showed intermediate values between the WT and *atg5*, and a similar trend was observed in *PpPINA* and *PpRSL4* expression levels. Thus, different extent of *PpPINA* inhibition in each mutant could explain differences in their IAA content, which in turn might result in distinct PpRSL4 expression levels. Previous work using *PpSHI1* over-expressing lines showed that higher expression levels led to a more pronounced senescence phenotype (Eklund et al., 2010), supporting the idea that different levels of auxin content and response might contribute to the phenotypic differences of *atg5* and *atg14* mutants.

Lastly, we demonstrate how autophagic flux is modulated across varying conditions of both carbon and nitrogen supply. Across a normal long-day cycle in complete nitrogen medium, autophagic flux was inhibited during the light periods and induced during the dark periods. Exposure to complete nitrogen deficiency resulted in a global reduction of autophagic flux, especially during the dark periods, apparently inverting the light-dark modulation seen in the complete medium. This inversion might be the result of the altered C/N supply ratio, resulting in autophagy being induced when this ratio is most unbalanced (either to C or N). In our previous work, we demonstrated the induction of autophagic flux in N deficiency under continuous light, i.e. a higher C/N supply ratio (Pettinari et al., 2022). In this work the added variation in carbon supply showed how both sources might interact in shifting the modulation of autophagic flux. However, exposure to external auxin (a strong promoter of caulonemal development) also decreased autophagic flux, which might suggest that conditions that favor the chloronemata to caulonemata transition could as well reduce autophagic activity in general. This explanation might initially seem inconsistent with our previous work, where we showed an increase in autophagic particles in both chloronemal and caulonemal apical and subapical cells under complete nitrogen deficiency (Pettinari et al., 2022). However, interestingly the number of autophagic particles in chloronemal apical cells was consistently higher than in caulonemal apical cells, independently of the conditions tested (Pettinari et al., 2022). Altogether, the evidence suggests that higher autophagic activity might be intrinsically associated with chloronemal cells and thus chloronemal-favouring conditions, whereas caulonemal-favouring conditions would decrease autophagy. Recent studies have demonstrated that autophagy is required to promote root hair longevity and pollen tube growth in vascular plants, which, like protonemata, depend on polar growth for their elongation (Feng et al., 2025; Yan et al., 2023). However, a key feature that sets apart moss protonemal cells from root hairs and pollen tubes is their capacity to continue dividing indeterminately throughout the moss life cycle, each one constituting a stem cell able to produce every cell type in the plant, ultimately generating three-dimensional structures and allowing the cycle to continue (Rounds & Bezanilla, 2013). In contrast to the growth of root hairs and pollen tubes, in protonemal cells, the autophagic process itself would probably not be enough to sustain growth under nutrient-limited conditions in the long term. Instead, the plant needs to change its developmental program and trigger the transition from the photosynthetically active chloronemata to the colonizing caulonemata. Thus, the determinate vs indeterminate growth of these tip-growing cells from these divergent plant lineages may reflect cell-type-specific autophagic functions as part of the mechanisms activated to cope with changing nutrient availability.

## CONCLUSION

In this work, we highlight autophagy as a key regulatory node that integrates changes to the C/N supply ratio with the developmental transition from chloronemata to caulonemata in *Physcomitrium patens*. We show how mutants deficient in the autophagic function display a loss of coordinated plasticity in response to a changing environment, showing an initially enhanced yet ultimately unsustainable response to nitrogen deficiency alongside an attenuated response to external sucrose. These mutants appear to have a heavily altered basal state (Figure 9B), evidenced by higher intrinsic caulonemal growth and modified internal sugar profiles even under non-stressful conditions. This shifted state is also reflected in internal auxin accumulation, differential auxin-related gene expression, and either diminished response or accelerated senescence upon auxin supplementation in the mutants. This altered auxin regulation and response likely provides a mechanistic link between the disrupted metabolic state of the mutants and their dysregulated caulonemal development. Alongside the previously described *atg5* and *atg7* mutants we describe the *atg14* mutant, which systematically showed an intermediate phenotype compared to *atg5* (including in auxin accumulation, gene expression, and responses to C/N availability), further supporting the link between autophagy, auxin, and the C/N ratio. Finally, we demonstrate how autophagic flux is dynamically regulated by the C/N supply ratio, being activated when this ratio is most unbalanced. We propose that high autophagic activity is associated with the chloroplast-rich chloronemata, whereas caulonemata-inducing conditions globally decrease autophagic flux (Figure 9A). This cell-specific autophagic regulation might reflect both the differing functions of these two cell types as well as the indeterminate nature of their growth. In summary, our findings suggest that upon alterations to environmental nutrient supply, autophagy acts as a key regulator in the system modulating an ordered and adaptive response, a capacity compromised in *atg* mutants (Figure 9). In doing so, it coordinates caulonemal development and growth with both external (C/N supply) and internal (auxin-related) signals.

**Figure 9.**
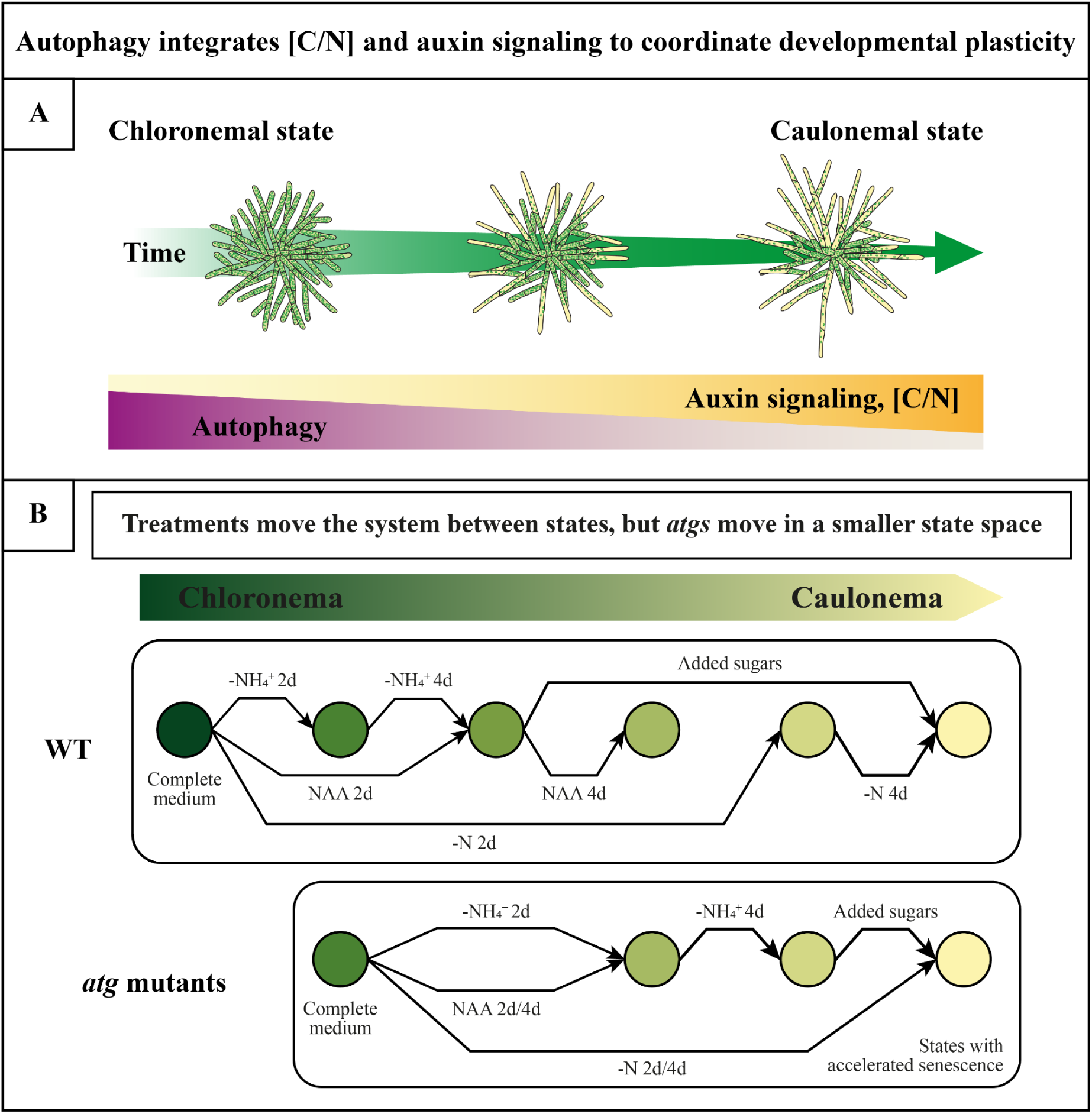
Conceptual model integrating the impact of autophagy during the chloronema-to-caulonema transition. A) During protonemal development, the system progresses from a chloronema-enriched state to a caulonema-enriched state over time. This transition correlates with increased auxin signaling and a higher [C/N] ratio. Autophagic activity, on the other hand, declines, consistent with its role as a regulatory node coordinating the transition. B) Along the chloronema-to-caulonema developmental axis (left-to-right, green-to-yellow), the system can occupy a range of developmental states. Under control conditions (complete medium), the WT plants occupy a highly chloronemal state. Different environmental or hormonal treatments can shift the system towards distinct caulonema-enriched states. In contrast, *atg* mutants display a basally shifted state even in complete medium, towards a more caulonemal state than the WT. Moreover, the number of possible states the *atg* system can access is reduced compared to the WT. Transitions between states are simplified: some treatments (e.g. -N) trigger a strong but ultimately unsustainable caulonemal response, whereas others (e.g. sugar supplementation) elicit a weaker effect. This results in a simpler, less adaptive network. Notably, *atg* mutants undergo accelerated senescence, particularly in more caulonemal states. Altogether, these differences suggest that *atg* plants operate within a smaller state space, reflecting lower developmental plasticity and adaptive capacity.

## Supporting information

List of primers used in this study

## Author contributions

G.P: Conceptualization, experimental design, performed experiments, formal analysis, and writing - original draft, Writing - review & editing. FL: Conceptualization, experimental design, performed experiments, formal analysis, and writing - original draft, Writing - review & editing. VM: Performed hormone quantification. MT: Performed hormone quantification. GR: Formal analysis, funding acquisition, writing - review & editing. MB: Methodology, supervision, writing - review & editing. RL: Conceptualization, experimental design, funding acquisition, supervision, project administration, validation, writing - review & editing. LS: Conceptualization, experimental design, performed experiments, formal analysis, funding acquisition, supervision, project administration, validation, writing - review & editing.

## Declaration of generative AI and AI-assisted technologies in the manuscript preparation process

During the preparation of this work the authors used DeepSeek in order to help readability and overall language, and to assist in code writing in R. After using this tool, the authors reviewed and edited the content as needed and take full responsibility for the content of the published article.

## Acknowledgments

We acknowledge CONICET for the Ph.D. scholarships to G.P and to F.L, and the Fulbright Commission and CFI Argentina for the FLB-CFI-2023-2024 research grant to G.P. We acknowledge Dr. Virginia Lobatto and Dr. Esteban Schenfield for technical support, and Dr. Damián Cambiagno and Dr. Ignacio Lescano for helpful discussions.

---

**Figure S1.**
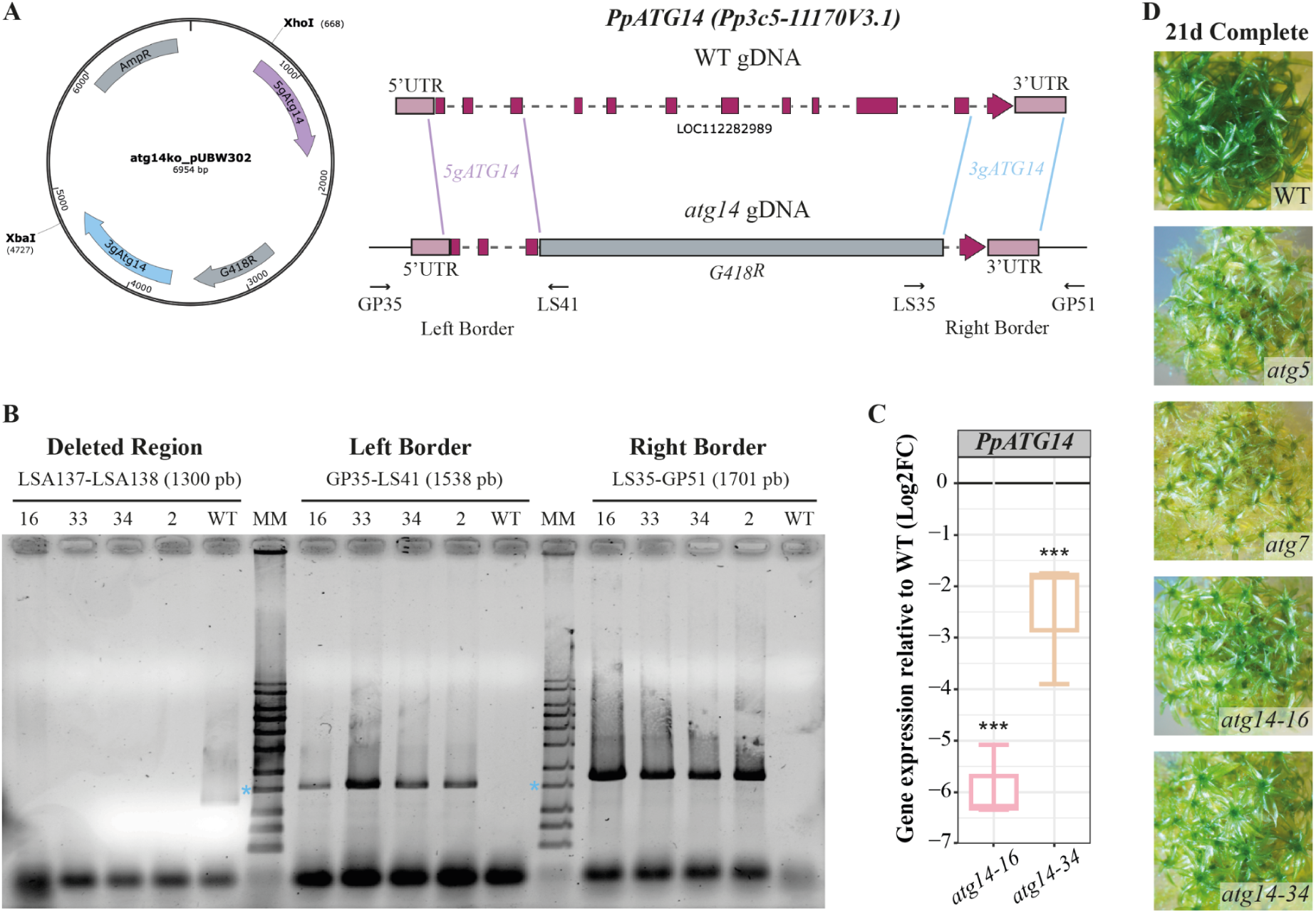
Generation and genotyping of *Ppatg14* mutant lines. A) Vector used for protoplast transformation, indicating the restriction enzymes used for linearization (left). Schematic representation of genomic DNA from WT plants and *atg14* mutants, showing the exon structure of PpATG14 and the replaced region (right). B) Genotyping of four independent *atg14* lines. Light blue asterisks indicate the 1500 bp marker band. C) Relative *PpATG14* transcript levels in *atg14-16* and *atg14-34* lines determined by RT–qPCR. Values are expressed relative to WT (black line) and represent mean ± s.d. of biological replicates (n = 3). ***p < 0.01 (Student’s *t*-test vs WT). D) Colony phenotype of WT, *atg5*, *atg7*, and two independent *atg14* lines (*atg14-16* and *atg14-34*) after 21 days of growth on complete medium.

